# DNA methylation regulates sex-biased gene expression in the house sparrow

**DOI:** 10.1101/2022.11.07.515394

**Authors:** Sarah L. Lundregan, Hannu Mäkinen, Heidi Viitaniemi, Bernt Rønning, Henrik Jensen, Arild Husby

**Author notes:** Senior author (Arild Husby).

## Abstract

Sexual dimorphism is often mediated by tissue-specific, differential gene expression, but the mechanisms that regulate these gene expression patterns are not well understood. Here, we investigate DNA methylation as a potential regulator of sex-biased gene expression in the house sparrow (*Passer domesticus*). First, we examine whether sex-bias in gene expression or DNA methylation is present in this species, and whether any sex differences are tissue-general or tissue-specific. Second, we assess the correlation between gene expression and DNA methylation at different genomic features in several tissues. Samples clustered by tissue type when looking at both gene expression and DNA methylation, and in gonads samples clustered according to sex. We demonstrate sex-bias in DNA methylation and gene expression on the Z-chromosome as well as on autosomes, but find that most of the sex-biased genes are tissue-specific and that the majority of sex-bias occurs in gonads, although some tissue-general sex differences were observed. This underscores the importance of choosing a tissue relevant to the studied phenotype in DNA methylation or gene expression studies. We find strong negative correlation between DNA methylation and gene expression at the transcription start site (TSS), and that the TSS of highly expressed genes is hypomethylated in comparison to the TSS of genes with low expression. Furthermore, we show that sex-biased DNA methylation can account for 14% of the sex differences in gene expression in gonads. Together these results indicate that DNA methylation differences between the sexes can provide a mechanistic explanation for sex-biased gene expression that likely contributes to trait sexual dimorphism in nature.

## Introduction

Sexual dimorphism is commonly observed in a broad range of traits in plants and animals, including morphology (Badyaev, 2002; Dunn et al.. 2001), physiology (Tarka et al., 2018; Gao et al., 2021; Vincze et al., 2022), behaviour (Lewis et al., 2002; Pellman et al., 2017; Carbia and Brown, 2020), and life history (Tarka et al., 2018; Lemaître et al., 2020). Sexual dimorphism occurs due to sex differences in the optimal phenotype that are established and maintained by selective processes, i.e. natural or sexual selection (Ellegren and Parsch, 2007; Harrison et al., 2015). Although sexual dimorphism is commonly observed, how such differences can evolve is a puzzle at the genetic level because females and males share the same genome apart from the sex chromosomes. The sex chromosomes are known to play an important role in sex-specific traits such as female fertility (Moghadam et al., 2012), female mate preference (Saetre et al., 2003), and male plumage traits (Saether et al., 2007), and sex chromosome linked gene expression is often strongly sex-biased in organisms such as birds that have incomplete dosage compensation (Dean and Mank., 2014). Nonetheless, in most species only relatively few genes on the sex chromosomes differ in copy number between females and males, which suggests that sex-biased expression of genes that are present in both sexes may be largely responsible for trait sexual dimorphism (Ellegren and Parsch, 2007; Mank et al., 2008). Differential gene expression between the sexes is often tissue-specific. While the most pronounced differences are commonly found in the gonads (see e.g. Harrison et al., 2015), somatic tissues also frequently exhibit high levels of sex-biased gene expression (Ellegren and Parsch 2007; Chatterjee et al., 2016; Bain et al., 2021), which may be a result of metabolic differences between females and males and may give rise to secondary sexual characteristics (Mank et al., 2008). The molecular mechanisms that regulate variation in gene expression are not well understood. Thus, exploring the molecular mechanisms that contribute to sex differences in phenotypic traits may enable us to better understand how these processes contribute to ecological and evolutionary dynamics in natural populations.

Sex-biased gene expression has been linked to phenotypic differences between the sexes in several vertebrate species. (Rinn and Snyder, 2005; Ellengren and Parsch 2007; Sharma et al., 2014), including in birds. Sex-biased gene expression in the collared flycatcher (*Ficedula albicollis*) is highly correlated with nucleotide diversity (Dutoit et al., 2018), which suggests that genetic diversity in natural populations may in part be maintained by sexual conflict. Large sex differences in chicken (*Gallus gallus*) behavior have been linked to sex-biased gene expression in the brain (Nätt et al., 2014). Non-random genomic distribution of genes with sex-biased expression has also been shown in chicken brain (Kaiser and Vera, 2006), but although genes with higher expression in males were over represented on the Z-chromosome many autosomal genes with sex-biased expression were identified. Similarly, extensive sex differences in brain gene expression have been demonstrated in common whitethroat (*Sylvia communis*) and in zebra finch (*Taeniopygia guttata*) and most genes identified were male-biased and Z-linked (Naurin et al., 2011). Male-biased expression of microRNAs encoded on the Z-chromosome that may be related to song behavior and neural plasticity has been demonstrated in zebra finch (Luo et al., 2012), and sex-biased expression of ribosomal proteins has also been linked to song system in this species (Tang and Wade, 2010). While sex differences in phenotype between individuals may be modulated by differences in gene expression, gene expression profiles of homologous tissues are generally highly conserved and vary less than gene expression profiles between different tissues from the same species (Sudmant et al., 2015). Such tissue-specific gene expression is well documented in birds, including chicken (Nie et al., 2010; Nätt et al., 2012; Wu et al., 2016; Bush et al., 2018), great tit (*Parus major*, Laine et al., 2019), flycatcher (*Ficedula*) species (Uebbing et al., 2016; Mugal et al., 2020), and tree swallow (*Tachycineta bicolor*, Bentz et al., 2019). Thus, sex-biased gene expression in birds may be expected to be highly tissue specific, and the most pronounced differences between the sexes are likely to be found in the gonads (Harrison et al., 2015), or on the Z-chromosome in somatic tissues (Kaiser and Vera 2006; Itoh et al., 2007).

DNA methylation is a well-characterised epigenetic modification that could regulate sex- and tissue-specific differences in gene expression (Hu and Barrett, 2017). DNA methylation is the addition of a methyl (-CH_3_) on the 5’ carbon of a cytosine residue to produce 5-methylcytosine. Cytosine methylation is most common at CpG sites in vertebrates, but may also be present at non-CpG sites, especially in brain tissue (Auclair and Weber, 2012). Cytosine methylation is often inversely related to gene expression when it occurs near the transcription start site (TSS), as it can inhibit transcription factor binding at the associated promoter or result in recruitment of chromatin modifying complexes (Deaton and Bird, 2011; Jones, 2012). DNA methylation patterns in birds appear similar to those in other vertebrates: methylation occurs mainly on cytosine residues in a CpG context, and aside from sites near the promoter region of genes much of the genome is methylated (Head, 2014). For example, in chicken cytosine methylation occurs predominantly at CpG sites, is higher in gene bodies, and in the TSS and promoter methylation is negatively correlated with gene expression (Li et al., 2011). The recently published great tit methylome demonstrated that DNA methylation levels dropped distinctly toward the TSS, and both CpG and non-CpG methylation at the TSS and promoter was negatively correlated with gene expression (Laine et al., 2016). Whereas a positive correlation between gene expression and gene body methylation has been demonstrated in many mammalian tissues (Jones, 1999), a negative correlation has been demonstrated in great tit brain tissue (Derks et al., 2016), which suggests that the relationship between non-TSS DNA methylation and gene expression may be tissue dependent (see also Jones, 2012).

DNA methylation at specific CpG sites is often tissue-specific in vertebrates and may regulate differential gene expression and biological functions of distinct tissue types (Deaton et al., 2011; Auclair and Weber, 2012). Tissue-specific expression of DNA methyltransferases has been found in duck (*Anas platyrhynchos domestica*, Yan et al., 2015) and zebra finch (Mishra et al., 2020). Tissue-specific DNA methylation has been demonstrated in chicken (Xu et al., 2007; Li et al., 2011; Nätt et al., 2012) and great tit (Derks et al., 2016; Watson et al., 2021), and DNA methylation at the promoter region of genes was negatively correlated with gene expression in both species (Li et al., 2011; Derks et al., 2016; Watson et al., 2021). Conversely, tissue-general temporal changes in DNA methylation have been demonstrated in the great tit, but associations between these changes in DNA methylation and gene expression were tissue and genomic location dependent (Lindner et al., 2021). DNA methylation could contribute to phenotypic differences between the sexes through regulation of sex-biased gene expression patterns, and DNA methylation patterns on autosomes as well as on the sex chromosomes can differ between females and males (McCarthey et al, 2014; Hu and Barrett, 2017; McCormick et al., 2017). In starlings (*Lamprotornis superbus*) unpredictable early-life environmental conditions may lead to sex-biased DNA methylation changes that impact fitness (Rubenstein et al., 2016). Furthermore, DNA methylation in the promoter region of specific genes has been negatively related to sex-biased gene expression in chicken, although differences in gene expression between females and males did not coincide with differences in promoter methylation on a genome-wide scale (Nätt et al., 2014).

There are few studies to date that have evaluated both sex-biased gene expression and DNA methylation patterns across several tissues (but see Luo et al., 2012; Bain et al., 2021), and studies that investigate these processes in natural populations are needed to broaden understanding of their effects on sex differences between individuals in nature. To identify tissue-general and tissue-specific sex-biased gene expression and determine the role of DNA methylation in this process, we analysed the methylomes of blood, brain, liver, and gonad, and the transcriptomes of brain, liver, and gonad from eight male and eight female wild house sparrows using RNA-seq and reduced representation bisulfite sequencing (RRBS). The aims of this study were three-fold. First, we analysed whether gene expression and DNA methylation were tissue-general or tissue-specific and whether there existed any sex-bias in gene expression or DNA methylation. Second, we investigated the relationship between gene expression and DNA methylation at different genomic features. Third, we assessed to what extent DNA methylation in blood can be used as a proxy for DNA methylation patterns in other more inaccessible tissues.

More specifically, we first examined genome-wide gene expression and DNA methylation patterns to determine whether similarity between samples was best explained by tissue or sex. Then we performed differential gene expression analyses and differential methylated region analyses in each tissue to identify tissue-specific and tissue-general, sex-biased, differentially expressed genes (DEGs) and differentially methylated regions (DMRs). Subsequently, we examined DNA methylation levels at six genomic features and looked at DNA methylation patterns close to the TSS and promoter in more detail. Lastly, we evaluated the correlation between gene expression and DNA methylation at four of the above genomic features as well as in the region ± 5 Kbp from annotated gene starts, and defined any overlap between genes that were differentially expressed and differentially methylated.

## Methods

### Study site and sampling

Sixteen house sparrows (eight females and eight males, aged from one to three years old) were sampled on the Island of Leka (65°5’N, 11°7’E) in Trøndelag, mid Norway. Leka is part of a long-term house sparrow study system and birds were sampled in connection with an artificial selection experiment on basal metabolic rate (BMR) that was conducted between 2012-2015 (Rønning et al., 2016). Data collection was carried out under permission from the animal experimentation administration (FOTS) of the Norwegian Food Safety Authority (NFSA), and in accordance with permits from the Norwegian Ringing Centre at Stavanger Museum and the Norwegian Environment Agency. Birds were captured by mist-netting within 10 days prior to the tissue sampling date (see Table 1 for individual sampling dates), and a 25 μl blood sample was taken from the brachial vein and stored in 96% ethanol for genomic DNA extraction and RRBS sequencing. On 16/02/2014, the 16 birds were collected using a blow to the cranium, and the brain, liver, and gonads were dissected immediately. Each organ was removed from surrounding tissue, weighed to the nearest 0.1 mg, then wrapped in aluminium foil and labeled with individual ring number and tissue type before freezing in liquid nitrogen. Time between death and collection of the final tissue sample for each individual was less than 12 minutes. Samples were subsequently stored at −80°C until further analysis. As the birds were collected in late winter, they were either in a non-breeding state or in a transition phase into the breeding state.

**Table 1:** Study sampling design. ID, sample date, sample location, individual sex, and age are shown. Both gene expression and DNA methylation sequencing data are available for brain, liver, and gonad tissue for all individuals. Only DNA methylation data is available for blood. For full details on number of sequenced reads and mapping success for each individual for the RNA-seq and RRBS data, see supplementary Table S1.

| ID | Location | Sample date blood | Sample date brain, liver, and gonad | Sex | Age |
| --- | --- | --- | --- | --- | --- |
| 8M71323 | Leka | 09/02/2014 | 16/02/2014 | Male | 3 |
| 8N06837 |  | 09/02/2014 | 16/02/2014 |  | 2 |
| 8N72694 |  | 11/02/2014 | 16/02/2014 |  | 2 |
| 8N73488 |  | 14/02/2014 | 16/02/2014 |  | 1 |
| 8N73907 |  | 11/02/2014 | 16/02/2014 |  | 1 |
| 8N87513 |  | 06/02/2014 | 16/02/2014 |  | 1 |
| 8N87539 |  | 07/02/2014 | 16/02/2014 |  | 1 |
| 8N87590 |  | 11/02/2014 | 16/02/2014 |  | 2 |
| 8N72676 | Leka | 07/02/2014 | 16/02/2014 | Female | 2 |
| 8N73920 |  | 11/02/2014 | 16/02/2014 |  | 1 |
| 8N87534 |  | 06/02/2014 | 16/02/2014 |  | 1 |
| 8N87536 |  | 06/02/2014 | 16/02/2014 |  | 1 |
| 8N87543 |  | 07/02/2014 | 16/02/2014 |  | 2 |
| 8N87582 |  | 11/02/2014 | 16/02/2014 |  | 2 |
| 8N87594 |  | 11/02/2014 | 16/02/2014 |  | 2 |
| 8N87595 |  | 11/02/2014 | 16/02/2014 |  | 2 |

**Table 2:** **A)** Number of DEGs found for each tissue in differential expression analysis using DeSeq2, and mean expression level ± SE of these genes for females and males separately. **B)** Number of DMRs found for each tissue in differential methylated region analyses using Methylkit, and mean methylation percentage ± SE at these regions for females and males separately.

| A) Differential expression analysis |  |  |  |
| --- | --- | --- | --- |
| Tissue | Number of DEGs | Mean expression females | Mean expression males |
| Brain | 300 | 7.377 ± 0.120 | 7.521 ± 0.121 |
| Liver | 45 | 6.977 ± 0.477 | 7.316 ± 0.542 |
| Gonad | 6838 | 6.735 ± 0.042 | 6.644 ± 0.041 |
| B) Differential methylated region analysis |  |  |  |
| Tissue | Number of DMRs | Methylation % females | Methylation % males |
| Blood | 413 | 39.90 ± 0.860 | 44.26 ± 0.966 |
| Brain | 465 | 46.43 ± 0.856 | 48.43 ± 0.901 |
| Liver | 454 | 45.57 ± 0.865 | 45.386 ± 0.904 |
| Gonad | 6820 | 41.52 ± 0.213 | 33.58 ± 0.255 |

### RNA extraction and sequencing

RNA was isolated from brain, liver, and gonad, but not blood samples. Each tissue sample (excluding a section taken for gDNA extraction) was weighed, then homogenized in ice-cold Trizol before isolation using the Zymo Direct-zol RNA Miniprep kit (Zymo Research) according to manufacturer instructions. Total RNA was eluted in 50 μl of nuclease free water and quality was checked using nanodrop before quantification using the Qubit RNA HS assay kit (ThermoFisher). The library preparation and sequencing were performed at the Genomics Core Facility (Department of Clinical and Molecular Medicine, Norwegian University of Science and Technology, Norway). Libraries were made using the Lexogen SENSE mRNA kit (Lexogen, Vienna) according to manufacturer instructions. Sequencing was done on one HiSeq 4000 flowcell (paired-end, 2x 150bp), and because samples were uniquely indexed they were pooled together and this pool was divided onto the eight lanes of the flow cell. Standard quality control with Fastqc was performed for the sequencing reads. Illumina adapter contamination was detected on both reads and was removed using Cutadapt with default parameters before alignment of reads to the *Passer domesticus* reference genome v 1.0 (Elgvin et al., 2017) using STAR aligner (Dobin et al., 2013). First an indexed genome was prepared using the genomic feature file containing intron-exon boundaries for the house sparrow gene models. Then each library was aligned to the reference genome with default parameters and the read count for each gene was counted. On average we obtained 54.8 (23.9 – 88.4) million raw reads per sequencing library. After quality filtering on average 36.9 (15.1 – 53.6) million reads were recovered per library. On average 81% (67.2 – 90) of the reads were uniquely aligned against the house sparrow genome. 18% of the unmapped reads were too short (< 50 bp) and the remaining unmapped reads were aligned to multiple regions. All the downstream analyses are based on the uniquely mapped reads. Altogether we obtained count data for 13004 genes for which the variance in counts between individuals across all tissues was larger than zero.

### Genomic DNA extraction and sequencing

An approximately 20 mg tissue section was excised from each frozen tissue sample for total genomic DNA (gDNA) extraction, after which the sample was immediately returned to −80°C to preserve RNA. Each tissue sample was lysed in proteinase K for 7 hrs at 56°C on a rocking platform. To obtain RNA free genomic DNA, RNase treatment was performed by adding 12 μl (10 mg/ml) RNaseA (ThermoFisher Scientific) to each sample lysate and incubating for 20 min at room temperature. The DNA extraction for brain, liver and gonad tissues was carried out using the Qiagen DNeasy Blood and Tissue Kit (Qiagen, USA) following manufacturer’s instructions. For blood samples, lysis was first carried out by incubation in Lairds buffer with 90 μg of proteinase K for 3 hours at 50°C. Automated extraction of gDNA from the lysate was carried out on a Biomek NX^p^ robot (Beckman Coulter, Miami, FL, USA) using the ReliaPrep Large Volume HT gDNA Isolation System (Promega, Madison, WI, USA). gDNA from tissue and blood samples was eluted in AE Buffer, and quality was assessed on a 1% agarose gel before quantification using the Qubit dsDNA HS Assay Kit (Invitrogen). Library preparation and subsequent RRBS sequencing were carried out at SciLife (Uppsala University, Sweden). Libraries were prepared according to Illumina protocol with some modifications. Briefly, samples were digested using the restriction enzyme MspI and the resulting DNA fragments were bisulfite treated then end repaired with DNA polymerase I, and A-overhangs were added to the 3’ ends of each fragment for adapter ligation. Individual sample libraries were barcoded using standard Illumina adapters and purified with Ampure XP beads (Beckman Coulter) The libraries were size-selected for insert sizes of 20-200 bp and quantified by qPCR. Six to seven libraries were pooled into the same sequencing lane. Each pool was sequenced 100 bp single-end on two flow cells of a HiSeq2500 sequencer using the HiSeq SBS sequencing kit version 4 (Illumina). Sequencing was conducted in two separate HiSeq runs to yield enough coverage per sample and later combined. An internal positive control (PhiX) was used to balance nucleotide diversity.

Before read alignment, low quality bases and adaptor contamination were trimmed using TrimGalore! 0.6.7 in -rrbs mode with default parameters. Trimmed sequencing reads were aligned against an in-silico RRBS digested, bisulfite converted version of the *Passer domesticus* reference genome v 1.0 (Elgvin et al., 2017) using BiSulfite Bolt aligner (Farrell et al., 2021) in single-end mode with default alignment parameters. Mean mapping efficiency was 86% (see Table S1 for individual sample details). Methylation bias plots (Hansen et al., 2012) for each sequenced individual were produced using MethylDackel v 0.3.0 to assess methylation percentage in each position of sequence reads, and 4 bp were subsequently trimmed from both the ‘5 and ‘3 end of reads. CpG sites were then identified for each sample using the “CallMethylation” function in BiSulfite Bolt, and mean methylation percentage across tissues was 27% (see Table S1 for individual sample details).

### Genome-wide gene expression and DNA methylation patterns

Prior to all analyses on the RNA-seq data, transcripts with low expression (counts less than 10 across all samples within each tissue) were removed to ensure high confidence. To determine whether sex was a good indicator of similarity in gene expression in brain, liver, and gonad, we performed distance based hierarchical clustering and PCA on the regularized log transformation procedure (rld) transformed gene expression values produced using DeSeq2 v3.14 (Love et al., 2014). To compare methylation patterns between tissues, only sites with a minimum of 5x coverage (including methylated and unmethylated counts) were used (313701 CpG sites, see Table S2). To determine whether sex predicted similarity in DNA methylation levels in blood, brain, liver, and gonad, we performed hierarchical Ward D2 clustering with Manhattan distance using the “cluster samples” function from the R package Methylkit (Akalin et al., 2012) and did principal component analysis (PCA) using the “PCASamples” function. We also calculated mean methylation at these sites for each tissue.

### Sex-biased differential expression

Differential expression of genes between the sexes was analysed separately for each tissue using the standard DeSeq2 protocol with age as a covariate in the full model. Benjamini-Hochberg correction of *p*-values was used and genes with an FDR *q*-value less than 0.05 were considered to be differentially expressed. Heat maps of the top 30 differentially expressed genes in each tissue were generated using the rld transformed expression values and the Pheatmap R package. Volcano plots of log2 fold change *q*-values were produced using the R package EnhancedVolcano. Functional analysis of DEGs was conducted using the Cytoscape plugin StringApp v1.7.0 (Doncheva et al., 2019) with a default confidence of 0.4 to construct a separate protein-protein interaction network (PPI) for each tissue. For gonad only differentially expressed genes that were also DMGs were selected for functional analysis to reduce the many functional terms to those that were most relevant. Subsequently, Weighted gene correlation network analysis (WGCNA, Langfelder and Horvath, 2008) was performed on the gonad data normalised counts to create modules of co-expressed genes. Correlation of the module eigengenes with sex was then calculated to identify gene modules that may be important for sexual dimorphism. Briefly, we created an unsigned network using a soft thresholding power of 4 (chosen using the “pickSoftThreshold” function with a cut-off value of 0.8), minimum module size of 20 transcripts, and a merge cut height of 0.25 to allow us to combine the similar modules from the same node into larger modules, before correlating the module eigengenes with sex. We then analysed hub genes (Pearson correlation for Module Membership > 0.8, *p*-value for the significance of the relationship with sex < 0.05) from sex correlated modules by using pathway analysis. Pathway analyses were conducted using StringApp, using the default confidence of 0.4 to construct a separate protein-protein interaction network (PPI) for each module. Subsequently “real hub” genes were identified for each network using a connectivity degree that selected the most connected ~5% of nodes on each network and functional analysis was performed on these “real hub” genes using StringApp.

### Sex-biased DNA methylation

Differential methylated region analysis was carried out using Methylkit, and was done separately for each tissue using sites with 5x coverage that were identified across all samples within each tissue (this resulted in 533416 CpG sites for analysis in blood, 644517 sites for brain, 852190 sites for liver, and 585205 sites for gonad, see Table S2). 5x coverage was used because this level of coverage has been shown to be sufficient when performing DMR analysis using Methylkit with several samples per condition (Piao et al., 2021). The genome was first tiled into windows of 300 bp using the “tileMethylCounts” function with step size 300 and a minimum of 5 CpG sites in each window. A window size of 300 bp was chosen because CpG sites within a few hundred base pairs of each other have been shown to exhibit greater spatial autocorrelation in DNA methylation levels than sites situated further apart (Eckhardt et al., 2006). Differential methylation analysis was then performed on these windows using the “calculateDiffMeth” function to implement logistic regression, with age as a covariate. Benjamini-Hochberg correction of *p*-values was used, and regions with a false discovery rate (FDR) *q*-value of less than 0.05 and with greater than 10% methylation difference between the sexes were considered to be differentially methylated. Differentially methylated genes (DMGs) were defined as genes with DMRs in their TSS or promoter. Functional analysis of DMGs was done on a per-tissue basis using StringApp with the parameters defined above. For gonad only DMGs that were also differentially expressed were selected for functional analysis, and to further reduce gonad associated functional terms to the most relevant terms, only “real hub genes” (the 5% of genes in the network with the highest degree connectivity) were selected. All functional analyses in this study were performed using human GO Biological Process, Reactome Pathway, Wiki Pathway, KEGG Pathway, and TISSUE annotations, and a redundancy cutoff of 0.6 was used to collapse redundant terms. Any terms with an FDR *q*-value less than 0.05 were considered to be enriched.

### Relationship between DNA methylation and gene expression

To investigate methylation patterns at different genomic features within each tissue, we used sites with 5x coverage that were identified across all samples within each tissue (see Table S2). The R package “GenomicFeatures” (Lawrence et al., 2013) was used to extract start and end positions for gene bodies (defined as the annotated gene start to annotated gene end, excluding the transcription start site), TSS (defined as the region 300 bp upstream to 50 bp downstream of the annotated gene start), promoters (defined as the region 3 Kbp to 301 bp upstream of the annotated gene start), and the region 10 Kbp to 3 Kbp upstream of the annotated gene start. BedTools v2.29.2 (Quinlan and Hall, 2010) was then used to assign CpG sites to these features so that methylation patterns in these different genomic locations could be compared. Repetitive elements were identified using the Early Grey transposable element (TE) annotation pipeline (Baril et al., 2021). Briefly, the *Aves* library was first used to annotate known repeats, then novel TEs were identified and refined using an iterative implementation of the “BLAST, Extract, Extend” process to characterize elements along their entire length (Platt et al., 2016). Lastly, overlapping and fragmented annotations were resolved using Earl Grey before final TE quantification. We assigned CpG sites to TEs including long-interspersed nuclear elements of type Chicken Repeat 1 (CR1s) and long terminal repeats (LTRs) to determine methylation levels at these TEs. To obtain detailed information on DNA methylation patterns near the TSS and promoter, regions ± 5 Kbp from annotated gene starts were divided into 40 consecutive tiles of 250 bp that were centred on the TSS, then mean number of CpG sites and mean methylation percentage at each tile was calculated.

To determine how DNA methylation influences gene expression in different tissues, Spearman’s rank correlation between mean DNA methylation and rld transformed gene expression was calculated. This was done on a per-tissue basis for each genomic feature (10 Kbp upstream, promoter, TSS, and gene body), and for each 250 bp tile in the region ± 5 Kbp from annotated gene starts (40 windows in total). Subsequently we grouped genes into low, mid, and high expression within each tissue (low = lowest expressed 20% of genes, high = highest expressed 20% of genes, mid = remainder of genes) and plotted this against mean methylation levels in the region ± 5 Kbp from annotated gene starts to assess whether methylation levels near the TSS were lower for the most highly expressed genes. We also evaluated any association between methylation difference between the sexes (male=1 and female=2) at DMRs in the TSS, promoter, or gene body and log2 fold change in gene expression at the corresponding genes. DMRs from differential methylated region analysis in Methylkit (regions with a *q*-value < 0.05 and methylation difference between the sexes of at least 10%) were plotted against corresponding genes with a log2 fold change greater than 0.5 using separate quadrant plots for each tissue. There are 4 possible quadrants for association between methylation level and gene expression: Q1 represents lower methylation in females leading to increased gene expression in females, Q2 represents higher methylation in females leading to increased gene expression in females, Q3 represents higher methylation in females leading to reduced gene expression in females, and Q4 represents lower methylation in females leading to reduced gene expression in females. For DMRs overlapping the TSS of genes we expect points to be over-represented in Q1 and Q3 compared to Q2 and Q4. For the gonad, Fisher exact tests were used to determine whether DMRs overlapping the TSS or promoter of genes were enriched in quadrants Q1 or Q3 compared to Q2 or Q4, whereby we compared the proportion of associations in quadrants Q1 and Q3 between the DMRs overlapping the TSS or promoter of genes and DMRs overlapping the gene body. Fisher exact tests were not done for brain or liver due to low number of DMRs with a log2 fold change greater than 0.5.

## Results

### Genome-wide gene expression and DNA methylation patterns

In the gene expression data, both hierarchical clustering (Fig. 1A) and PCA (Fig. 1B) showed that samples clustered according to tissue type, and in the gonad samples clustered according to sex. Difference in mean methylation between the sexes was highest in the gonads, and low in blood, brain, and liver tissue (Fig. 1C, *p*-values from unpaired *t*-Tests between the sexes were 0.005 for gonad, 0.609 for blood, 0.797 for brain, and 0.617 for liver). Samples clustered according to tissue in PCA on these shared sites, and in the gonad samples clustered according to sex (Fig. 1D). A similar pattern was observed in hierarchical Ward D2 clustering with Manhattan distance (Fig. S1). Thus, similar patterns of clustering were detected in the gene expression data and DNA methylation data.

**Figure 1:**
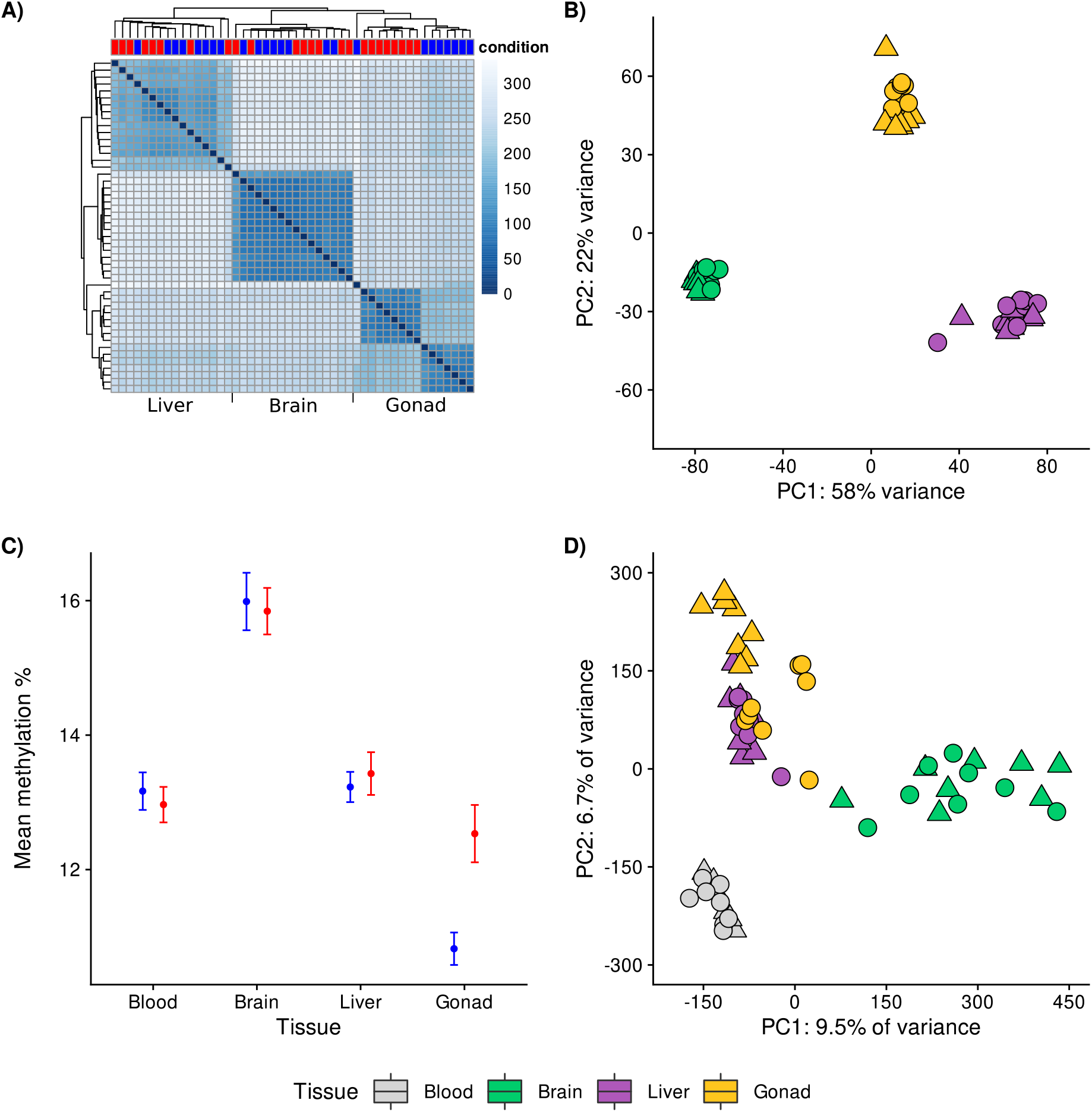
Summary analyses of RRBS and RNA-seq data. **A)** Distance between samples using the rld transformed gene expression data, sample distance was lowest between samples within tissues except for in the gonad where sex (female = red, male = blue) also predicted sample distance. **B)** Principal component analysis of the rld transformed gene expression data. Points are coloured by tissue type (see legend) and shape indicates sex (female = circle, male = triangle). **C)** Comparison of mean methylation percentage in blood, brain, liver and gonad using the 313701 sites that had 5x coverage in all samples. Females are shown in red and males are shown in blue. Error bars represent standard error of the mean methylation percentage between individuals. **D)** Principal component analysis of methylation profiles based on the same 313701 sites with 5x coverage in all samples. Points are coloured by tissue type (see legend) and shape indicates sex (female = circles, male = triangles).

### Sex-biased differential expression

Unsurprisingly, gonad tissue had the highest number of transcripts that showed evidence of differential expression between the sexes (Figures S2 and S3) and differentially expressed genes were distributed across the genome (Fig. 2A). 6838 DEGs were detected in gonad, 300 in brain, and 45 in liver (see Table S3 for an overview of all differentially expressed genes for each tissue). The majority of the DEGs in brain and liver were located on the sex (Z) chromosome and were more highly expressed in males (Fig. 2A), which is likely because males are the homogametic sex in birds and due to the lack of an overall dosage compensation in birds (Itoh et al., 2007; Uebbing et al., 2013). Most DEGs detected in gonad were tissue-specific, however, 63% of brain DEGs were also differentially expressed in gonad, and 38% of liver DEGs were also differentially expressed in gonad. Only ~2% of all DEGs were tissue-general (Fig. 2B), these tissue-general DEGs were all located on the Z-chromosome and showed male-biased expression in all tissues. Subsequently, separate functional analyses were carried out on all DEGs for brain and liver (Table S4, Figure S4A). For brain 17 functional terms were enriched, and no functional terms were enriched for liver. 22 functional terms were enriched for gonad when using only differentially expressed genes with DMRs in their TSS or promoter.

**Figure 2:**
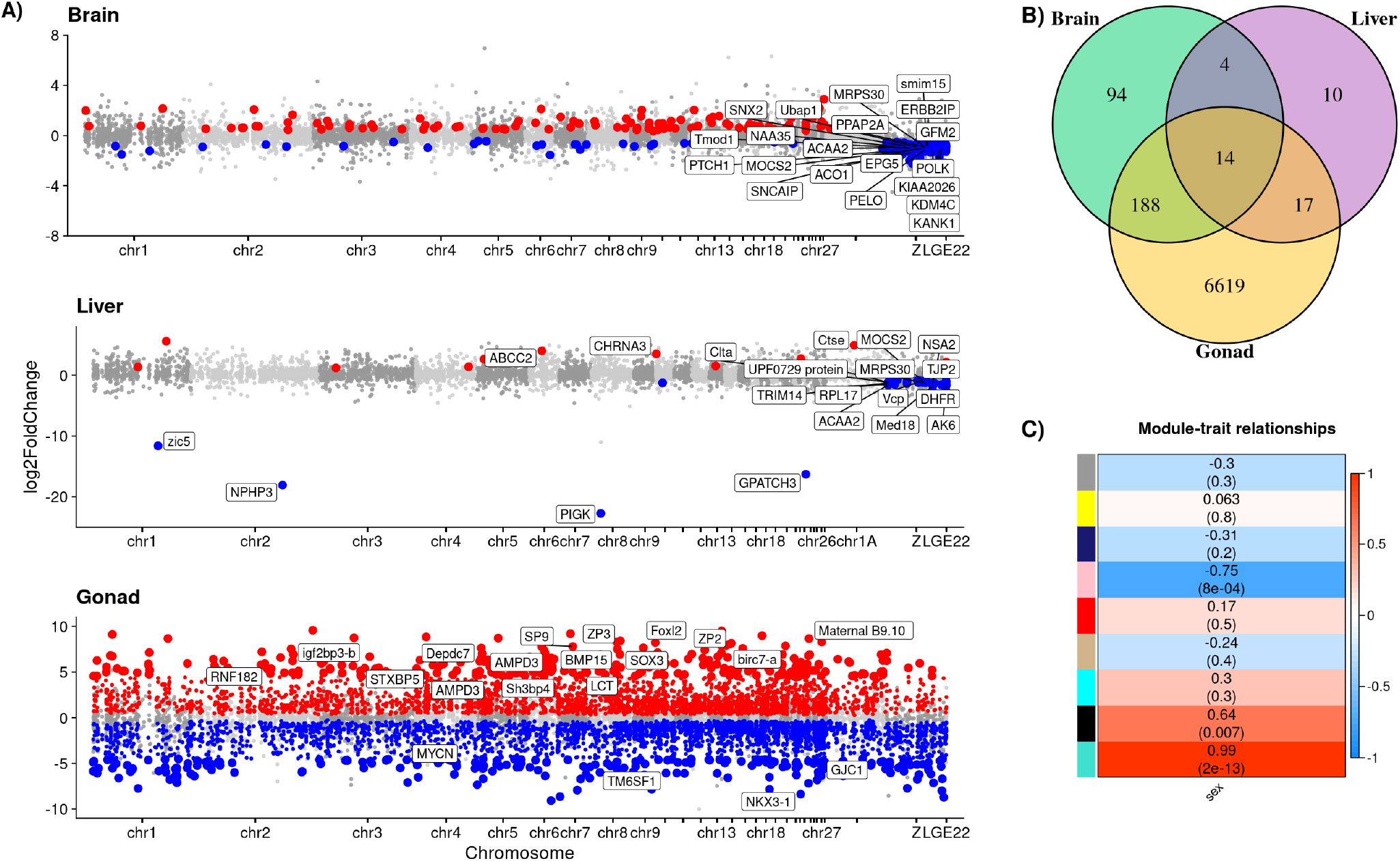
Results from analysis of the gene expression data. **A)** Manhattan plots of DEG analysis results with log2 fold change plotted against chromosomal location of genes. DEGs in the TSS or promoter of genes are coloured according to methylation difference between females and males (DEGs with higher expression in females are shown in red, and DEGs with higher expression in males are shown in blue). The 20 genes with the lowest *q*-value are labelled for each tissue. **B)** Venn diagram showing tissue-specific and tissue-general differentially expressed genes (DEGs). The majority of gonad DEGs were tissue-specific, however, 63% of brain DMGs, and 38% of liver DMGs were also differentially expressed in gonad. Only ~2% genes were tissue-general and were differentially expressed in all tissues. **C)** WGCNA analysis results. 9 modules were identified, three of which were correlated with sex. When male = 1 and female = 2, the turquoise module was positively correlated with sex (r = 0.99, *p* = 2e^−13^), as was the black module (r = 0.64, 0.007), whereas the pink module was negatively correlated with sex (r = −0.75, *p* = 8e^−4^).

Patterns of co-expression among transcripts in the gonad were investigated using WGCNA (see supplementary Figures S5-S7 for parameter and clustering details). Three modules showed strong evidence of correlation with sex: the turquoise (r = 0.99, *p* = 2 × 10^−13^) and black (r = 0.64, *p* = 0.007) modules were positively correlated with sex and thus contained a majority of genes that were more highly expressed in females, whilst the pink module was negatively correlated with sex (r = −0.75, *p* = 0.001) and contained more genes that were highly expressed in males (Fig. 2C, Fig. S8). Details of the transcripts belonging to the modules are provided in Table S5. Based on a combination of strong co-expression module membership, high significance of relationship with sex, and high PPI network connectivity, 135 “real” hub genes were determined for the turquoise module, 14 “real” hub genes were identified for the black module, and 33 “real” hub genes were identified for the pink module. PPI networks consisting of the “real” hub genes from each sex-associated module were highly interconnected (Fig. S8). 539 functional terms were enriched for the turquoise module that was positively correlated with sex (Table S6, Fig. S8A), 47 functional terms were enriched for the pink module that was negatively correlated with sex (Table S6, Fig. S8B), and 22 functional terms were enriched for the black module that was positively correlated with sex (Table S6, Fig. S8C). Any functional terms that may be relevant to phenotypic differences between the sexes are discussed in detail below.

### Sex-biased differential methylation

As expected, gonads had the highest number of genes that showed evidence of differential methylation between the sexes (Fig. 3A, Table S7). Of the 42275 tiles analysed (300 bp tiles with at least 5 CpGs), 6820 DMRs were detected in gonad, 1165 of which were located in the TSS or promoter of 760 differentially methylated genes. In blood, 36683 tiles were analysed and 413 DMRs were detected, 79 of which were in the TSS or promoter of 63 differentially methylated genes. In brain, 47301 tiles were analysed and 465 DMRs were detected, 67 of which were in the TSS or promoter of 61 differentially methylated genes, and in liver 53728 tiles were analysed and 454 DMRs were detected, 60 of which were in the TSS or promoter of 53 differentially methylated genes. For an overview of differentially methylated genes for each tissue, see Table S7, and see Fig. S9 for *q*-value distributions from DMR analysis on each tissue. Interestingly, sex-biased DMRs within transposable elements LTRs and CR1s were detected for each tissue (Fig. 3B). In gonad 74 of the 531 LTRs covered by the RRBS data had overlapping DMRs, and 92 of 725 CR1s covered by the RRBS data had overlapping DMRs. Most gonad differentially methylated genes were tissue-specific, however, 56% of blood DMGs were also differentially methylated in gonad, 51% of brain DMGs were also differentially methylated in gonad, and 60% of liver DMGs were also differentially methylated in gonad. Only ~0.1% of all DMGs were tissue-general and were differentially methylated in all tissues (Fig. 3C). Nonetheless, methylation levels at DMRs were similar between tissues, with the exception of comparisons with gonad (Fig. S10). Functional analysis was carried out separately for blood, brain, and liver using all genes with DMRs in the TSS or promoter (Table S4). No enrichment of functional terms was found for these tissues. 22 functional terms were identified for gonad in functional analysis using genes with DMRs in their TSS or promoter that were also differentially expressed (Table S4). Most of the enriched terms for gonad have been linked to sex differences in previous studies.

**Figure 3:**
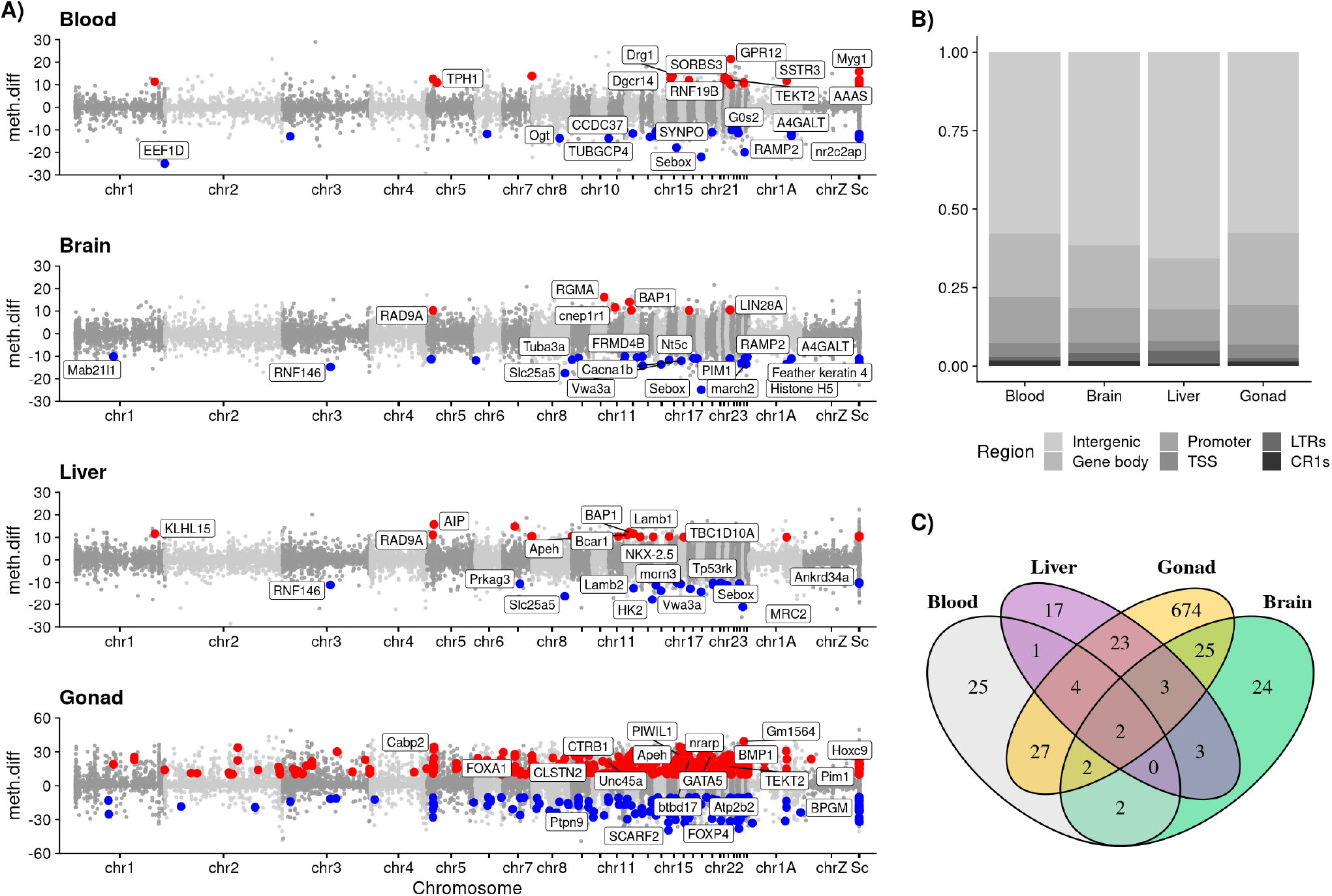
Differentially methylated region (DMR) analysis results. **A)** Manhattan plots of DMR analysis results. DMRs in the TSS or promoter of genes are coloured according to methylation difference between females and males (DMRs that had higher methylation levels in females are shown in red, and DMRs that had higher methylation levels in males are shown in blue). The 20 genes with the lowest *q*-value are labelled for each tissue. **B)** Proportion of DMRs assigned to different genomic features for each tissue. **C)** Venn diagram showing tissue-specific and tissue-general differentially methylated genes (DMGs) that had DMRs in the TSS or promoter. The majority of gonad DMGs were tissue-specific, however, 56% of blood DMGs, 51% of brain DMGs, and 60% of liver DMGs were also differentially methylated in gonad. Only 2 genes were tissue-general i.e. differentially methylated in all tissues.

### Relationship between DNA methylation and gene expression

Methylation levels were lowest at the TSS and promoter of genes, higher in the 10 Kbp region upstream of annotated gene starts, and highest in gene bodies. Transposable elements had relatively high methylation levels, and methylation was higher in LTRs than CR1s (Fig. 4A). There was very strong evidence that methylation levels differed between all genomic features (pairwise Wilcoxon rank sum tests, see Table S8). Closer evaluation of methylation levels in the region ± 5 Kbp from annotated gene starts demonstrated that mean methylation percentage was lowest at the TSS and promoter of genes and increased until ~2.5 Kbp from the annotated gene start (Fig. 4B). Number of CpGs was higher in the TSS and promoter of genes so there were more 250 bp tiles containing at least 1 CpG in the windows closest to the TSS (Fig. 4B), this is expected because CpG islands are predominantly found near the TSS or promoter of genes. There was strong evidence that mean DNA methylation level was negatively correlated with gene expression for all analysed genomic features in brain, liver, and gonad (Table S9). Spearman’s rank correlation between DNA methylation and gene expression was highest at the TSS, high at the promoter of genes, and lowest in the gene body and 10 Kbp upstream region (Fig. 4C, Fig. S11). Further evaluation of the correlation between DNA methylation and gene expression at the region ± 5 Kbp from annotated gene starts (Fig. 4D) showed that the negative correlation between DNA methylation and gene expression is strongest at the TSS (r = −0.361 in brain, r = −0.396 in liver, and r = −0.446 in gonad) and reaches a minimum by ± 3 Kbp from the annotated gene start. Within tissues, DNA methylation levels close to the TSS showed the sharpest decrease for the 20% of genes with the highest expression levels, but remained unchanged for the 20% of genes with the lowest expression levels (Fig. 4E).

**Figure 4:**
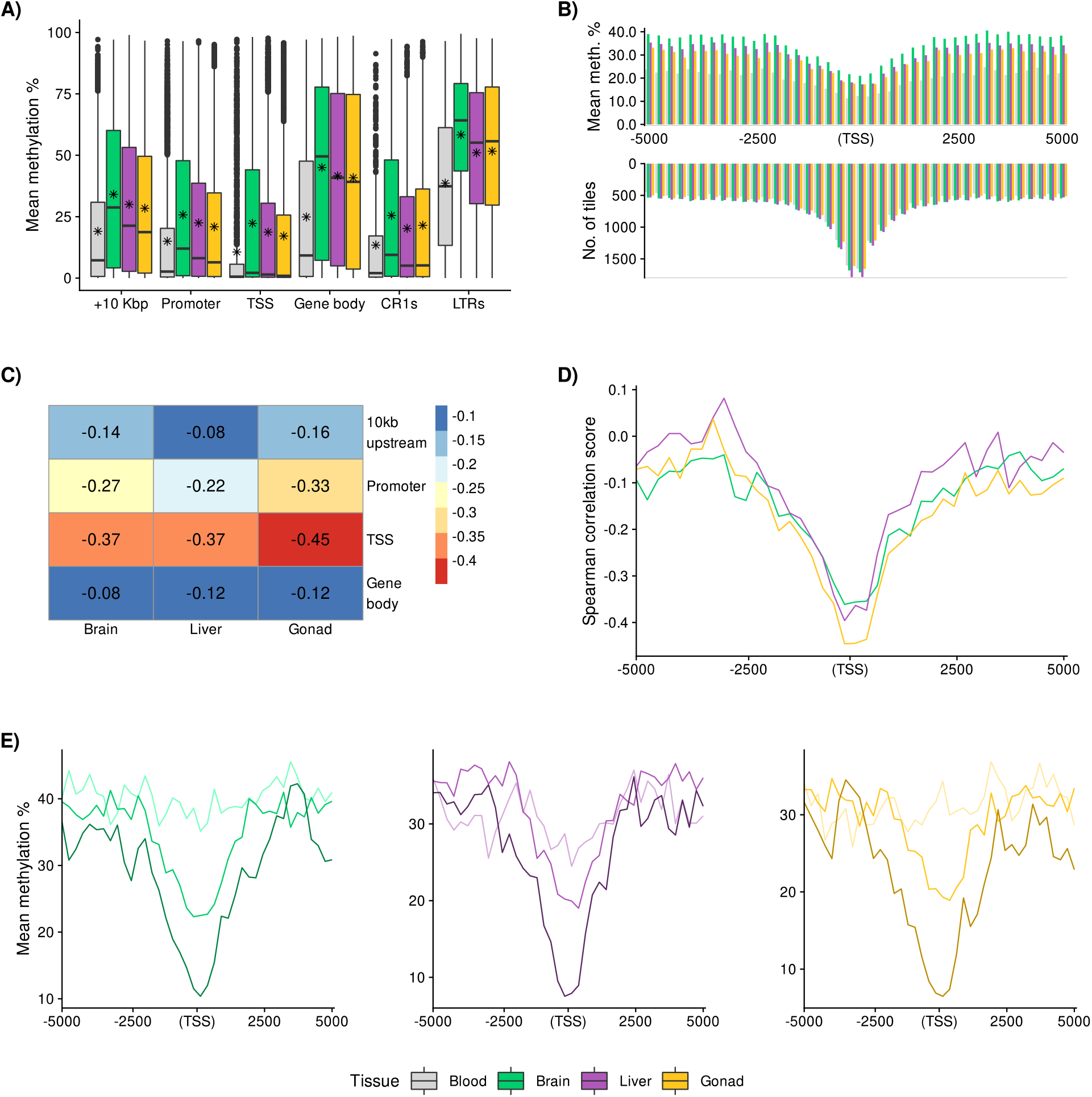
**A)** Boxplot of mean methylation percentage in different genomic features using sites that had 5x coverage for all samples within each tissue. Boxes are coloured according to tissue type (see legend), black asterisks represent mean methylation percentage, and the horizontal black line represents the median methylation percentage. There was very strong evidence that all distributions were different from each other (*p* = 1.381e^−4^ for the comparison between promoters and CR1s, and *p* < 7.518e^−29^ for all other comparisons) based on results of pairwise Wilcoxon rank sum tests (see Table S3). **B)** Comparison of number of tiles and mean methylation percentage at these tiles in the region ± 5 Kbp from annotated gene starts, using sites that had 5x coverage for all samples within each tissue. The relationship between mean DNA methylation level at distinct genomic features and rld transformed gene expression. **C)** Spearman’s rank correlation between DNA methylation level and gene expression at the 10 Kbp upstream region, promoter, TSS, and gene body. Gene expression is negatively correlated with DNA methylation at all genomic features, and this negative correlation is strongest at the TSS of genes. **D)** Spearman’s rank correlation between gene expression and DNA methylation ± 5 Kbp from annotated gene starts. The negative correlation between gene expression and DNA methylation is strongest at the TSS (r ~ 0.4) and reaches a minimum of r ~ 0.1 around ± 3 Kbp from the annotated gene start. **E)** Mean methylation percentage ± 5 Kbp from annotated gene starts for genes with high, medium, and low expression (for each tissue higher gene expression is indicated by darker lines). DNA methylation at the TSS is lowest for the highest expressed 20% of genes, and DNA methylation does not decrease closer to the TSS for the lowest expressed 20% of genes.

A similar proportion of DEGs had at least one 300 bp tile containing at least five CpG sites within the TSS or promoter in brain (30%), liver (33%), and gonad (38%). The relationship between sex-biased differential methylation of genes and sex-biased differential expression of genes was most consistent in the gonad. 47.9% of genes that were differentially methylated in gonad showed evidence of differential expression between the sexes, and 14% of gonad DEGs with methylation information had DMRs in their TSS or promoter (Fig. 5A, Table. S10). By contrast, in brain only 2% of DEGs with methylation information had DMRs in their TSS or promoter, and in liver 0% of DEGs with methylation information had DMRs in their TSS or promoter (Table S10). To further investigate the relationship between changes in DNA methylation and changes in gene expression we produced quadrant plots using DMRs in the gene body, TSS, or promoter of genes that had a log2 fold change between females and males of at least 0.5 at the corresponding genes (Fig. 5B, Fig. S12). For the gonad 903 DMRs passed this fold change threshold, but for brain and liver there were few DMRs that passed the threshold (for brain 14 DMRs and for liver 27 DMRs). In gonad, genes with DMRs overlapping the TSS were overrepresented within the expected quadrants (Q1 and Q3) when compared to genes with DMRs in the gene body (Fig. 5B, Fisher’s Exact Test *p* = 0.021). In contrast, genes with DMRs overlapping the promoter were not overrepresented in Q1 and Q3 (Fishers Exact Test *p* = 0.871). For brain and liver, the genes with DMRs in their TSS or promoter were randomly distributed across all quadrants, although number of data points for both tissues was low (Fig. S12).

**Figure 5:**
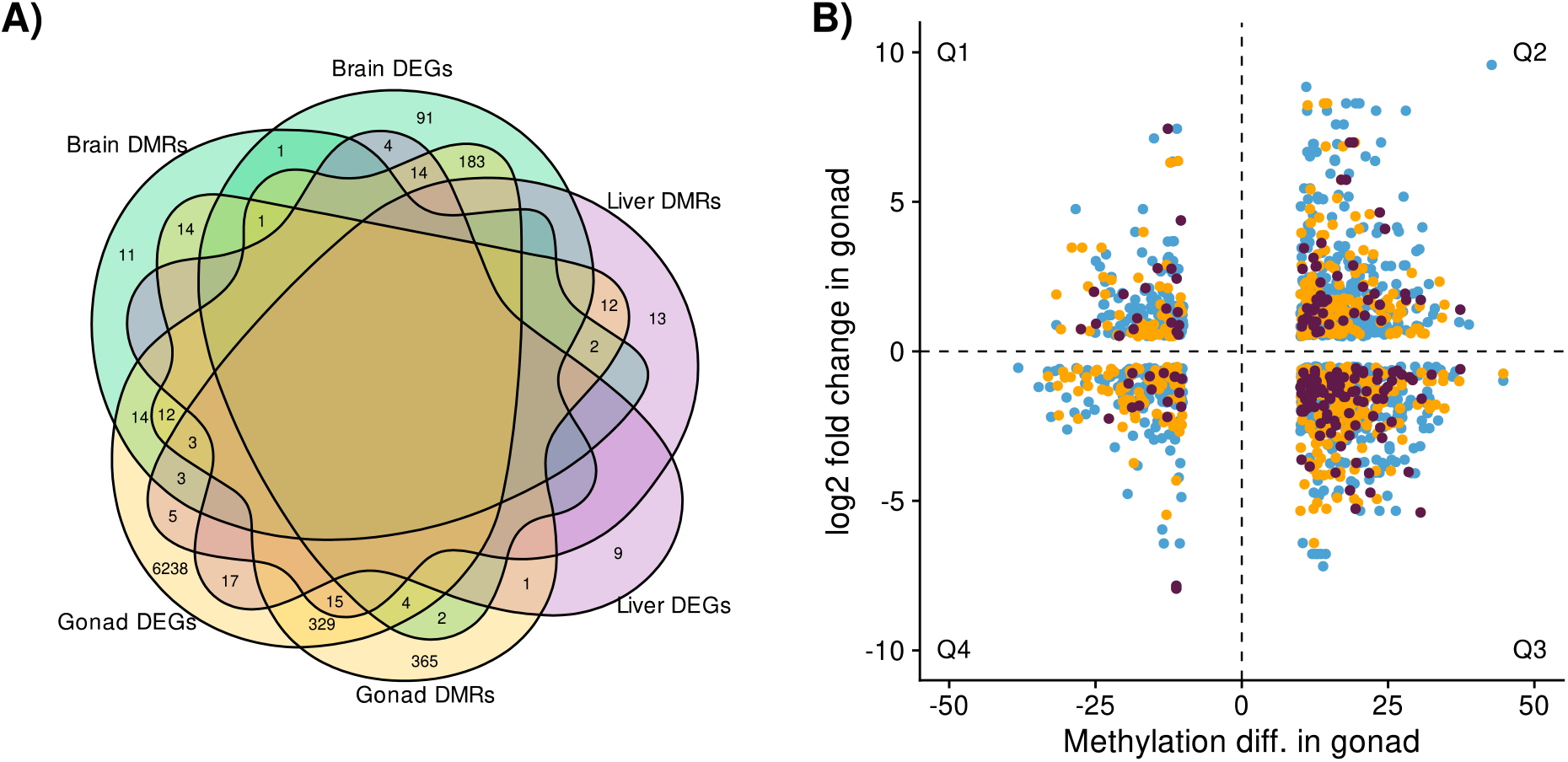
**A)** Venn diagram showing the relationship between sex-biased differential methylation of genes and sex-biased differential expression of genes. This relationship was strongest in the gonad, where 47.9% of genes with DMRs in the TSS or promoter were also DEGs. By contrast, in brain only 3.3% of DMRs were also DEGs, and in liver 0% of DMRs were also DEGs. However, 54.1% of genes that were differentially methylated in the brain were differentially expressed in gonad, and 49.1% of genes that were differentially methylated in liver were differentially expressed in gonad. **B)** Methylation difference between females and males of DMRs in the gonad in relation to log2 fold change in gene expression between females and males in the gonad. DMRs in the promoter of genes are shown in orange, DMRs in the TSS are shown in dark purple, and DMRs in the gene body are shown in blue. The 4 quadrants are separated by dotted lines and labelled Q1-Q4. Over representation of points in Q1 and Q3 is expected for DMRs located in transcription start sites.

## Discussion

Although gene expression and DNA methylation differences between tissues are well documented (see e.g Li et al., 2011; Nätt et al., 2012, Derks et al., 2016; Laine et al., 2019; Lindner et al., 2021) and several studies have demonstrated sex-biased gene expression or DNA methylation (see e.g. Nätt et al., 2014; Harrison et al., 2015; Rubenstein et al., 2016), there are few studies to date that evaluate both sex-biased gene expression and DNA methylation patterns across several tissues (but see Luo et al., 2012; Bain et al., 2021). Here, we demonstrate sex-biased gene expression on autosomes as well as the Z-chromosome, and identify sex-biased DMRs genome-wide. We found that most DEGs and DMRs were tissue-specific and that sex-biased gene expression and DNA methylation were most apparent in the gonad, although some tissue-general sex differences were observed. Furthermore, we expand on current knowledge by showing that sex-biased DNA methylation can account for approximately 14% of the sex differences in expression in gonad. Thus, DNA methylation differences between the sexes are likely to contribute to sex-biased gene expression and sex differences in phenotypes in the house sparrow.

### Genome-wide gene expression and DNA methylation patterns

Tissue of origin explained most of the similarity in gene expression between samples and in gonad samples also clustered according to sex (Figures 1A and 1B). Difference in mean methylation between the sexes was most apparent in the gonads (Figure 1C), and results of hierarchical clustering and PCA indicated that each tissue had distinct DNA methylation patterns (Figures 1D and S1). Thus samples clustered similarly in principal component analyses on both the DNA methylation and gene expression data. Unique transcriptome patterns of different tissues have previously been demonstrated in wild collared (*Ficedula albicollis*) and pied flycatchers (*Ficedula hypoleuca;* Uebbing et al., 2016), great tits (Laine et al., 2019), and tree swallows (Bentz et al., 2019). Taken together, our results support previous findings that DNA methylation and gene expression may be tissue-specific in wild birds.

### Tissue-specific and tissue-general sex differences

A large majority of sex-biased differentially expressed genes were detected in gonad, which is as expected. However, DEGs were also detected in brain, and liver, and approximately 50% of these DEGs were also differentially expressed in gonad. Thus, while differential expression in somatic tissues predicted differential expression of the same genes in the gonad to some degree, most DEGs were gonad-specific (Fig. 2B). Similarly, blood-derived measurements of genome-wide DNA methylation did not reflect those obtained from gonad (Figures 1C, and 4A), and whilst approximately 50% of the DMGs identified in somatic tissues were also differentially methylated in gonad, the majority of DMGs were gonad specific (Figure 3C). The lack of overlap between sex-biased genes in the gonad and the relatively few sex-biased genes identified in somatic tissues underscores the importance of choosing a tissue appropriate to the studied phenotype when carrying out differential methylation or gene expression analyses (Husby et al., 2020). This is in contrast to previous work that showed that temporal DNA methylation changes in red blood cells were tissue-general in the great tit (Lindner et al., 2021), therefore, blood derived values may still be a suitable proxy for methylation in less accessible tissues for some phenotypes and the suitability of blood should be assessed on a study to study basis.

### Sex-biased genes

Gonad tissue had the greatest number genes that were differentially expressed between the sexes, followed by brain, a pattern which has also been demonstrated in chicken (Mank et al., 2008). Of the DEGs we identified in somatic tissues (brain and liver), the vast majority were located on the Z-chromosome and had higher expression in males (Fig. 2A). Dosage compensation is less efficient in birds than in mammals, and higher expression of Z-linked genes has been demonstrated in male collared flycatcher (Uebbing et al., 2013), zebra finches and chickens (Itoh et al., 2007). However, both studies found considerable variation in the male-to-female ratio of gene expression among genes which suggests that partial dosage compensation may have evolved on a per-gene basis in birds (Itoh et al., 2007; Uebbing et al., 2013). Our findings support the idea that Z-linked genes are more likely to control sexual differentiation in somatic tissues in birds. Interestingly, 75% of the Z-linked genes that were DEGs in brain were among the 30% most highly expressed genes on the Z chromosome, and for liver 96% of Z-linked DEGs were among the 30% most highly expressed genes on Z. This is in agreement with previous findings that the most highly expressed genes on avian Z-chromosomes are most likely to display male-biased expression (Naurin et al., 2012), a phenomenon which suggests male-driven selection on Z-linked loci. Several functional terms were associated with brain DEGs: the top TISSUE terms “internal female genital organ”, “uterus”, “endocrine gland”, and “embryonic structure” were all related to reproductive function. The KEGG Pathway “ribosome biogenesis in eukaryotes” was identified, and various ribosomal genes have been linked to sexual differentiation in the brain related to song-learning in zebra finch (Tang and Wade, 2010; Qi et al., 2012; Acharya and Veney, 2013). The GO Biological Processes “nitrogen compound metabolic process” was important, and uric acid (a product of nitrogen metabolism) is a potent antioxidant that is found at higher levels in female birds (Hartman et al., 2006). No functional terms were associated with liver DEGs.

In accordance with the results from differential expression analyses, sex-biased differential methylation was highest in gonad, although DMGs were also identified in blood, brain, and liver (Figure 3A). Sex-biased DNA methylation often occurs on autosomes in addition to the sex chromosomes (McCarthy et al., 2014; McCormick et al., 2017), and here DMGs were distributed throughout the genome for all tissues. Nonetheless, no enriched functional terms were identified for blood, brain or liver DMGs (Table S4), and methylation differences at these DMGs rarely translated to sex-biased gene expression (only 3.3% of brain DMGs were also DEGs, and no liver DMGs were also DEGs). Conversely, 50% of gonad DMGs were also DEGs, and several relevant functional terms were identified for the subset of gonad DMGs that were also differentially expressed (Table S4). The highly relevant TISSUE term “female reproductive gland” was detected, and genes associated with the “digestive gland” term were linked to pancreatic function and insulin secretion, both of which are known to display sex differences (Hall et al., 2014). Reactome Pathway terms “selenoamino acid metabolism” and “meiotic synapsis” were identified and extensive sexual dimorphism in these processes has previously been demonstrated (Seale et al., 2018; Cahoon and Libuda, 2019). Furthermore, “estrogen receptor-mediated signalling” is central to female reproductive function (Fuentes and Silveyra, 2019), so the observed sex-biased methylation of genes involved in this pathway points to involvement of DNA methylation in reproductive processes in the house sparrow. GO Biological Process terms relating to hormone response and signalling were also identified. Steroid hormones are integral to normal follicular development and function, and have been implicated in hormone-mediated maternal effects in birds (Groothuis and Schwabl, 2008). Lastly, “ribosomal large subunit assembly” was identified. Z-linked ribosomal large subunit (RPL) genes have been linked to sexual dimorphism in the zebra finch song system (Tang and Wade, 2010), and both RPL and ribosomal small subunit (RPS) genes show greater expression in male songbirds (Qi et al., 2012; Acharya and Veney, 2013).

In the present study, several sex-biased differentially methylated TEs were identified, and the majority of these TEs were found in the gonads where the largest overlap between DMGs and DEGs was observed (12 sex-biased TEs were identified in blood, 19 in brain, 22 in liver, and 163 in gonad). Thus, sex-biased methylation of TEs may contribute to divergent TE activity in gonads between house sparrow females and males and, correspondingly, to phenotypic sex differences in this species. Differentially methylated TEs were located throughout the genome in all tissues, which is in contrast to previous studies that demonstrate that the avian W chromosome contains over half of potentially active CR1s (Dechaud et al., 2019; Peona et al., 2021). However, it is important to note that any TEs that are only present on the W chromosome are not covered by this dataset because the W chromosome is not included in the house sparrow reference genome (Elgvin et al., 2017), and because TEs need to be present in the genomes of both sexes in order to assess sex-bias in methylation levels.

### Sex-correlated co-expression modules

Several functional terms of interest were identified by examining the “real hub” genes that were most highly correlated with sex in the WGCNA analysis on the gene expression data (Table S6). For the turquoise module that was positively correlated with sex (when male = 1 and female = 2), the tissue terms with the lowest FDR values: “digestive gland”, “internal female genital organ”, “cervical carcinoma cell”, “male reproductive system”, and “fetus”, were all reproductive tissues. Many GO Biological Process terms relating to large and small ribosomal subunit biogenesis and assembly were again enriched. Several enriched Reactome Pathway terms were related to metabolism, as were the GO Biological Process terms “macromolecule metabolic process” and “cellular macromolecule biosynthetic process”, and the KEGG Pathways “insulin signalling pathway”, “thyroid hormone signalling pathway” and the “FoxO signalling pathway” that regulates glucose and lipid metabolism (Gross et al., 2008). This enrichment for metabolism-related terms may indicate large metabolic differences between females and males. Such sex differences in metabolism have previously been demonstrated in model organisms (Cui et al., 2021; Gao et al., 2021; Huidobro et al., 2021), as well as in the house sparrow (Rønning et al., 2016). The relevant terms “growth hormone synthesis, secretion and action” and “progesterone-mediated oocyte maturation” were also identified using KEGG Pathway. Vascular endothelial growth factor A (VEGFA) is a critical regulator of sex-specific vascular development in the testes (Sargent et al., 2015) and the Wiki Pathway term “VEGFA-VEGFR2 signalling pathway” was enriched. The Wiki Pathway term “EGF/EGFR signalling pathway” was also enriched, and EGF controls several female reproductive processes including LH signal transmission in ovulation (Richani and Gilchrist., 2018) and endometrial function during pregnancy (Large et al., 2014). Several functional terms relating to regulation of reactive oxygen species (ROS) were enriched. ROS levels influence a multitude of female reproductive processes including ovulation and maintenance of pregnancy (Rizzo et al., 2012), and sex differences in oxidative physiology have been demonstrated in wild birds (Vincze et al., 2022). Finally, many functional terms that were enriched for the turquoise module related to immunity, which suggests that sex-differences in immune processes may exist in the house sparrow, a phenomenon that has recently been demonstrated several species of wild birds (Valdebenito et al., 2021; Vincze et al., 2022). For the pink module that was negatively correlated with sex, many of the enriched functional terms related to ciliary motility and ciliary transport (the top term was the Reactome Pathway term “intraflagellar transport”, and the GO Biological Process terms “protein localization to cilium”, “Regulation of intraciliary retrograde transport”, “intraciliary retrograde transport”, and “Protein localization to non-motile cilium” were identified). Intraflagellar transport is important for spermatogenesis and male fertility (San-Agustin et al., 2015), and both motile and non-motile cilia are integral to male reproductive function (Girardet et al., 2019). Furthermore, proper regulation of ROS is critical for normal sperm function and male fertility (Mannucci et al., 2022), and the GO Biological Process terms “regulation of reactive oxygen species biosynthetic process” and “regulation of peroxidase activity” were enriched. The MAPK and RANKL/RANK pathways were identified using Wiki Pathway, and both pathways are known to regulate male reproductive function (Li et al., 2009; Blomberg Jensen et al., 2021). Interestingly, the Reactome Pathway term “EPH-ephrin signalling” was enriched, and ephrin signalling has been shown to influence aggression in male mice (Sheleg et al., 2015). Although the majority of terms identified using pink module “real hub” genes were unique to the pink module and were linked to male fertility, the “VEGFA-VEGFR2 signalling pathway” was also enriched for the turquoise module, as was the term “infectious disease”. Conversely, the TISSUE terms identified using “real hub” genes from the black module were the same as for the turquoise module, and many of the functional terms that were enriched for the black module overlapped with those that were enriched for the turquoise module: “ribonucleoprotein complex assembly” and “formation of the 43S complex” were important, as were “constitutive signalling by EGFRvIII”, “positive regulation of catabolic process”, and “influenza Viral RNA Transcription and Replication”. However, “inositol phosphate metabolism” (KEGG Pathway) that is important for insulin signalling (Hakim et al., 2012) was unique to the black module, as was the “RHOBTB2 GTPase cycle” (Reactome Pathway) that regulates vascular function and blood pressure (Loirand and Pacaud., 2014). Sex differences in vascular function are well studied in model organisms (Pabbidi et al., 2018).

In addition to the large overlap between the turquoise and black co-expression modules from WGCNA analysis, many of the functional terms that were enriched for these co-expression modules overlap with the functional terms that were enriched for gonad using the intersect between DEGs and DMGs. As expected, several terms relating to steroid hormone-mediated signalling were enriched for gonad in differential methylation and gene expression analyses as well as WGCNA analysis, which indicates that primary sex hormone signalling is crucial for sexual differentiation in the house sparrow. Interestingly, several terms relating to amino acid metabolism and immunity were enriched in both analyses, which is in agreement with previous studies that demonstrate extensive sex differences in these processes (see e.g. Seale et al., 2018; Valdebenito et al., 2021). Sex-biased expression of ribosomal proteins has been previously been linked to song system in zebra finch (Tang and Wade, 2010; Qi et al., 2012; Acharya and Veney, 2013). Here, several terms relating to large and small ribosomal subunit assembly were enriched across analyses, thus sex-biased expression of RPL and RPS genes in gonad may be central to sexual dimorphism in the house sparrow (Table 3, Tables S4 and S6).

**Table 3:** Summary of gene co-expression modules identified for gonad using WGCNA. Only modules that were strongly correlated with sex are shown. Network/module is the number of genes found by STRING analysis out of the whole set of module genes that passed the selection threshold (ModuleMembership > 0.8 and treatment *p*-value < 0.05). The top 5 “real hub” genes for each module are shown and were chosen based on strong co-expression module membership, high significance of relationship with sex, and highest PPI network connectivity.

| Module colour | Number of genes | DEGs in module | Network/module | Real hub genes | Real hub gene symbol | Gene name |
| --- | --- | --- | --- | --- | --- | --- |
| Turquoise | 6768 | 5211 | 2838/3202 | 135 | RPS15A | 40S ribosomal protein S15a |
|  |  |  |  |  | RPL27A | 60S ribosomal protein L27a |
|  |  |  |  |  | RPL23A | 60S ribosomal protein L23a |
|  |  |  |  |  | RPL32 | 60S ribosomal protein L32 |
|  |  |  |  |  | RPL35A | 60S ribosomal protein L35a |
| Pink | 2659 | 1192 | 733/782 | 33 | HSPB11 | Intraflagellar transport protein 25 homolog |
|  |  |  |  |  | TTC30B | Tetratricopeptide repeat protein 30B |
|  |  |  |  |  | LRRK2 | Leucine-rich repeat serine/threonine-protein kinase 2 |
|  |  |  |  |  | WDR35 | WD repeat-containing protein 35 |
|  |  |  |  |  | IFT57 | Intraflagellar transport protein 57 homolog |
| Black | 1199 | 364 | 330/351 | 14 | RPL3 | 60S ribosomal protein L3 |
|  |  |  |  |  | RPS13 | 40S ribosomal protein S13 |
|  |  |  |  |  | RPL15 | 60S ribosomal protein L15 |
|  |  |  |  |  | RPL9 | 60S ribosomal protein L9 |
|  |  |  |  |  | RPS14 | 40S ribosomal protein S14 |

### Relationship between gene expression and DNA methylation

DNA methylation levels in the house sparrow were lowest at the TSS and promoter of genes, higher in the 10 Kbp region upstream of annotated gene starts, and highest in gene bodies (Figures 4A and 4B), which is similar to DNA methylation patterns found in other studies in wild passerines (Laine et al., 2016; Viitaniemi et al., 2019). Mean DNA methylation in blood was lower than for brain, liver, and gonad at all genomic features (Fig. 4A). Reduced methylation in blood in comparison with other tissues has previously been documented (Derks et al., 2016), and in Viitaniemi et al. (2019) mean methylation of great tit blood was ~14% which is similar to the mean blood methylation level of ~13% found here (Fig. 1C). Intragenic CpG islands are more likely to be methylated in brain (Maunakea et al., 2010), which could contribute to the high brain methylation levels seen here (Fig. 1C), and, as is the case for house sparrows in the present study, TEs have been shown to be hypermethylated in great tit brain compared to blood (Derks et al., 2016). Here, very high mean methylation levels were found at LTR transposable elements across all tissues (Fig. 4A). Such high methylation levels likely contribute to LTR silencing and are expected because many LTRs are endogenous retroviruses (ERVs) some of which have been shown to be active in the avian genome (Peona et al., 2021). Spurious transcription of LTRs can cause genomic instability, especially when it disrupts essential genes or regulatory regions (Dechaud et al., 2019). By contrast, methylation of CR1 TEs was comparatively low in all tissues (Fig. 4A), this may be because the majority of CR1s in the avian genome are truncated from their 5’ end so are unable to retrotranspose (Hillier et al., 2004). High methylation levels of LTRs (specifically ERVs) compared to CR1s (LINEs) has previously been demonstrated in mice (Elmer et al., 2021) and similar high methylation of TEs has been found in great tit (Derks et al., 2016).

We assessed the correlation between DNA methylation and gene expression at four of the above genomic features in the house sparrow genome (10 Kbp upstream of the start position of annotated genes, the gene’s promoter, TSS, and gene body), and found that the negative correlation between DNA methylation and gene expression was strongest at the TSS and promoter of genes in all analysed tissues (Fig. 4C). Consistent with these results, negative correlation between CpG methylation in the TSS and promoter region and gene expression has previously been demonstrated in the great tit (Derks et al., 2016; Laine et al., 2016) as well as in other vertebrates (Deaton and Bird, 2011). In addition, it has been shown that the relationship between gene body methylation and gene expression is tissue-specific and that increased gene-body methylation in brain correlates with increased transcription (Maunakea et al., 2010). Correspondingly, we observed that the correlation between gene body methylation and gene expression was less negative in brain tissue. Closer examination of the region ± 5 Kbp from the annotated gene start (Figures 4B and 4D) revealed that tiles in the region ± 500 bp from the gene start had the highest CpG densities but showed the strongest negative correlation with gene expression in all tissues. This is in agreement with previous analyses on the correlation between DNA methylation and gene expression near annotated gene starts (see e.g. Derks et al., 2016; Laine et al,. 2016; Luo et al., 2018), and supports the notion that DNA methylation is more likely to inhibit transcription factor binding or result in recruitment of chromatin modifying complexes when it occurs at the TSS or promoter of genes (Deaton and Bird, 2011; Jones, 2012). The most highly expressed genes had the lowest methylation levels at the TSS across all tissues (Fig. 4E), which has similarly been shown in the great tit (Derks et al., 2016; Laine et al., 2016). The comparative hypermethylation of the TSS of the lowest expressed genes demonstrated in the present study suggests that DNA methylation plays an important role in gene silencing in the house sparrow.

Accordingly, in gonad approximately 14% of differentially expressed genes had DMRs in their TSS or promoter and 50% of DMGs showed sex-biased expression, which suggests that DNA methylation differences between the sexes may regulate sex-biased gene expression in the gonads. Although the proportion of DEGs that had corresponding methylation data was similar for all tissues (approximately 35%), only a small proportion of DEGs identified in brain or liver had DMRs in the TSS or promoter. This, in combination with our findings that most of the DEGs identified in brain and liver were male biased and located on the Z-chromosome, suggests that sex-biased methylation in somatic tissues is predominantly due to an overall lack of dosage compensation (Itoh et al., 2007; Dean and Mank 2014) in the house sparrow, and that factors other than DNA methylation may regulate the expression of sex-related autosomal genes in the somatic tissues of birds. Furthermore, the small overlap between gonad DEGs and DMGs and any sex-biased genes identified in somatic tissues demonstrates that sex-differences in gene expression and DNA methylation in the house sparrow are highly tissue-specific. Such tissue-specific differences in gene expression and DNA methylation may be directly related to tissue-specific functions (Husby, 2020). While we found little overlap between the functional terms found for brain DEGs and those found for gonad DEGs, terms relating to ribosome assembly were enriched in both brain and gonad. Thus, ribosome function, which has previously been linked to sex differences in the zebra finch song system (Tang and Wade 2010; Qi et al., 2012; Acharya and Veney, 2013), may contribute to sex differences in related behavioural traits in the house sparrow.

### Limitations and future directions

Birds in this study were sampled in connection with an artificial selection experiment on basal metabolic rate (BMR), which represents a potential source of bias. However, clustering analyses of the DNA methylation and gene expression data found no evidence that samples clustered according to BMR group (results not shown), and the few metabolism related functional terms that were identified have previously been linked to sex-differences in other species. Therefore, it is unlikely that this artificial selection experiment impacted our results. We compare methylation levels between blood, brain, liver and gonad, as well as examine whether blood derived measures of DNA methylation can be used as a proxy for methylation levels in other tissues. However, blood samples analysed in the current study were collected and stored in a different manner to the other analysed tissues (whole blood samples were stored in ethanol at −20°C, whereas tissue samples were immediately frozen in liquid nitrogen and stored at −80°C for subsequent RNA extraction). Previous studies have found that global DNA methylation may increase with storage time in frozen whole blood (Huang et al., 2017; Schroder and Steimer, 2018), whilst other studies that examined several different storage methods indicate that global DNA methylation profiles of whole blood are stable across a range of storage durations and conditions (Groen et al., 2018; Li et al., 2018). Nonetheless, differing storage conditions may result in small site-specific methylation changes (Groen et al., 2018). Therefore, although we do not expect the different storage of blood compared to other tissues in the current study to strongly influence results, the reported site-specific DNA methylation correlations between blood and the other analysed tissues may be downward biased. Furthermore, due to long-term storage at −20°C it was not possible to extract RNA from blood samples for gene expression analysis. Consequently, we cannot draw definite conclusions about the effects of DNA methylation in blood on gene expression, although we expect that the negative correlation between DNA methylation at the TSS and promoter of genes and gene expression that was observed in brain, liver, and gonad should also be observed in blood. Future work should explore the effect of blood DNA methylation on blood gene expression in the house sparrow, and use fresh whole blood samples to rule out any impacts of storage method on methylation levels.

We used an RRBS approach due to its cost effectiveness, which enabled us to sample more individuals. Although RRBS provides a sparse sampling of the genome compared to whole genome bisulfite sequencing (WGBS) data (Husby, 2022), RRBS sequencing data is enriched for CpG dense regions of the genome, such as the TSS and promoter of genes for which the correlation between DNA methylation and gene expression is most well established (Gu et al., 2011). Because DNA methylation levels exhibit spatial autocorrelation, sites situated close to each other often have more similar DNA methylation levels than sites situated further apart (Eckhardt et al., 2006). Consequently, DMRs may be of greater functional interest than single differentially methylated sites, however, the sparse sampling of RRBS data means that there will generally be less CpG sites in a region of given window size than for WGBS data (Lea et al., 2017). This represents a second drawback of the RBBS approach because comparatively few tiles with a given number of CpG sites will be present in a RRBS dataset. Nonetheless, in RRBS datasets a large proportion of these tiles are likely to be in CpG rich regulatory regions so a larger proportion of detected DMRs may be within genomic features that have a well categorised effect on gene expression.

We identified higher mean methylation levels of LTR transposable elements as compared to CR1s, and found evidence for sex-biased methylation of TEs in the gonad, as well as in somatic tissues. Comprehensive evaluation of the contribution of DNA methylation at TEs to sex differences in the house sparrow are beyond the scope of this study. However, future work should explore potential contribution of TE silencing to phenotypic differences between the sexes in the house sparrow by assessing the correlation between DNA methylation at TEs and TE expression in brain, liver, and gonad, and determine whether sex-biased differential methylation of TEs translates to differences in gene expression between the sexes. More generally, as technological advancements pave the way for larger epigenetic studies in natural populations, future work should aim to explore how DNA methylation and any corresponding differences in gene expression vary across environmental conditions, as well as control for confounding genetic effects (Husby, 2022). This will broaden understanding of how these processes contribute to ecological and evolutionary dynamics in nature.

## Conclusion

Here we evaluate both sex-biased gene expression and sex-biased DNA methylation patterns across several tissues in a natural population of house sparrows. We show distinct gene expression and DNA methylation patterns of four tissue types and demonstrate that there is a strong negative correlation between DNA methylation and gene expression at the TSS of genes in this species. Very low methylation levels were observed at the TSS of the most highly expressed genes whilst the TSS of genes with low expression were comparatively hypermethylated, which suggests that DNA methylation levels play a central role in governing gene expression in the house sparrow. Most sex-biased genes were tissue-specific, although some sex differences were also observed across multiple tissues, and this tissue-specific nature of sex-bias demonstrates that DNA methylation patterns in accessible tissues do not always predict DNA methylation and gene expression changes in target tissues. Accordingly, we find that sex-bias is strongest in gonad, and that sex-biased gene expression more often coincides with sex-biased DNA methylation in this tissue. Furthermore, functional analyses showed that sex-biased expression of ribosomal genes, as well as genes related to steroid hormone signalling, ciliary motility and intraciliary transport, and immunity, are likely to contribute to sex differences in the house sparrow. As such, this study broadens current knowledge on the contribution of DNA methylation dynamics to sexual dimorphism in wild birds.

## Acknowledgements

We would like to thank the inhabitants and especially the farmers on Leka, whose hospitality made this work possible. We also thank Randi Røsbak for her assistance with sample processing and RNA extraction. This study was funded by grants from the Research Council of Norway to H.J. (302619) and A.H. (239974), and by funding from the Norwegian University of Science and Technology to S.L.L. The empirical research was carried out in accordance with permits from the Norwegian Food Safety Authority and the Ringing Centre at Stavanger Museum, Norway. The authors acknowledge support from NTNU Genomics Core Facility and NTNU High Performance Computing Group. The authors also acknowledge support from the National Genomics Infrastructure in Stockholm funded by Science for Life Laboratory, the Knut and Alice Wallenberg Foundation and the Swedish Research Council, and SNIC/Uppsala Multidisciplinary Center for Advanced Computational Science for assistance with massively parallel sequencing and access to the UPPMAX computational infrastructure

## Author contributions

S.L.L, A.H, and H.J designed the study. B.R and H.J collected the field data and tissue samples. S.L.L extracted DNA and RNA for sequencing. S.L.L analysed the data with guidance from H.M, H.V, A.H and H.J. The article was written by S.L.L and H.V with input from all authors.

## Data availability

Phenotypic and sequence data used in this study will be made available on Dryad upon acceptance of this article.

## References

Acharya, K. D., & Veney, S. L. (2013). Sexually dimorphic expression and estradiol mediated up-regulation of a sex-linked ribosomal gene, RPS6, in the zebra finch brain. Developmental Neurobiology, 73(8), 599–608. https://doi.org/10.1002/dneu.22085

Akalin, A., Kormaksson, M., Li, S., Garrett-Bakelman, F. E., Figueroa, M. E., Melnick, A., & Mason, C. E. (2012). MethylKit: a comprehensive R package for the analysis of genome-wide DNA methylation profiles. Genome Biology, 13(10), R87. https://doi.org/10.1186/gb-2012-13-10-R87

Auclair, G., & Weber, M. (2012). Mechanisms of DNA methylation and demethylation in mammals. Biochimie, 94(11), 2202–2211. https://doi.org/10.1016/j.biochi.2012.05.016

Badyaev, A. V. (2002). Growing apart: an ontogenetic perspective on the evolution of sexual size dimorphism. Trends in Ecology & Evolution, 17(8), 369–378. https://doi.org/10.1016/S0169-5347(02)02569-7

Bain, S. A., Marshall, H., de la Filia, A. G., Laetsch, D. R., Husnik, F., & Ross, L. (2021). Sex-specific expression and DNA methylation in a species with extreme sexual dimorphism and paternal genome elimination. Molecular Ecology, 30(22), 5687–5703. https://doi.org/10.1111/mec.15842

Baril, T., Imrie, R., & Hayward, A. (2021). Earl Grey. https://doi.org/10.5281/zenodo.5654616

Bentz, A. B., Thomas, G. W. C., Rusch, D. B., & Rosvall, K. A. (2019). Tissue-specific expression profiles and positive selection analysis in the tree swallow (*Tachycineta bicolor*) using a de-novo transcriptome assembly. Scientific Reports, 9(1), 1–12. https://doi.org/10.1038/s41598-019-52312-4

Blomberg Jensen, M., Andreassen, C. H., Jørgensen, A., Nielsen, J. E., Juel Mortensen, L., Boisen, I. M., Schwarz, P., Toppari, J., Baron, R., Lanske, B., & Juul, A. (2021). RANKL regulates male reproductive function. Nature Communications, 12(1), 1–15. https://doi.org/10.1038/s41467-021-22734-8

Bush, S. J., Freem, L., MacCallum, A. J., O’Dell, J., Wu, C., Afrasiabi, C., Psifidi, A., Stevens, M. P., Smith, J., Summers, K. M., & Hume, D. A. (2018). Combination of novel and public RNA-seq datasets to generate an mRNA expression atlas for the domestic chicken. BMC Genomics, 19(1), 1–19. https://doi.org/10.1186/s12864-018-4972-7

Carbia, P. S., & Brown, C. (2020). Seasonal variation of sexually dimorphic spatial learning implicates mating system in the intertidal Cocos frillgoby (*Bathygobius cocosensis*). Animal Cognition, 23(4), 621–628. https://doi.org/10.1007/s10071-020-01366-3

Chatterjee, A., Lagisz, M., Rodger, E. J., Zhen, L., Stockwell, P. A., Duncan, E. J., Horsfield, J. A., Jeyakani, J., Mathavan, S., Ozaki, Y., & Nakagawa, S. (2016). Sex differences in DNA methylation and expression in zebrafish brain: a test of an extended ‘male sex drive’ hypothesis. Gene, 590(2), 307–316. https://doi.org/10.1016/j.gene.2016.05.042

Cui, L., Zhang, X., Cheng, R., Ansari, A. R., Elokil, A. A., Hu, Y., Chen, Y., Nafady, A. A., & Liu, H. (2021). Sex differences in growth performance are related to cecal microbiota in chicken. Microbial Pathogenesis, 150, 104710. https://doi.org/10.1016/j.micpath.2020.104710

Dean, R., & Mank, J. E. (2014). The role of sex chromosomes in sexual dimorphism: Discordance between molecular and phenotypic data. Journal of Evolutionary Biology, 27(7), 1443–1453. https://doi.org/10.1111/jeb.12345

Deaton, A., & Bird, A. (2011). CpG islands and the regulation of transcription. Genes & Development, 25(10), 1010–1022. https://doi.org/10.1101/gad.2037511.1010

Deaton, M., Webb, S., Kerr, A. R. W., Illingworth, R. S., Guy, J., Andrews, R., & Bird, A. (2011). Cell type – specific DNA methylation at intragenic CpG islands in the immune system. Genome Research, 21(7), 1074–1086. https://doi.org/10.1101/gr.118703.110.Freely

Dechaud, C., Volff, J. N., Schartl, M., & Naville, M. (2019). Sex and the TEs: Transposable elements in sexual development and function in animals. Mobile DNA, 10(1), 1–15. https://doi.org/10.1186/s13100-019-0185-0

Derks, M. F. L., Schachtschneider, K. M., Madsen, O., Schijlen, E., Verhoeven, K. J. F., & van Oers, K. (2016). Gene and transposable element methylation in great tit (*Parus major*) brain and blood. BMC Genomics, 17(1), 332. https://doi.org/10.1186/s12864-016-2653-y

Dobin, A., Davis, C. A., Schlesinger, F., Drenkow, J., Zaleski, C., Jha, S., Batut, P., Chaisson, M., & Gingeras, T. R. (2013). STAR: Ultrafast universal RNA-seq aligner. Bioinformatics, 29(1), 15–21. https://doi.org/10.1093/bioinformatics/bts635

Doncheva, N. T., Morris, J. H., Gorodkin, J., & Jensen, L. J. (2019). Cytoscape StringApp: Network analysis and visualization of proteomics data. Methods Molecular Biology, 18(2), 623–632. https://doi.org/10.1021/acs.jproteome.8b00702

Dunn, P. O., Whittingham, L. A., & Pitcher, T. E. (2001). Mating systems, sperm competition, and the evolution of sexual dimorphism in birds. Evolution, 55(1), 161–175. https://doi.org/10.1111/j.0014-3820.2001.tb01281.x

Dutoit, L., Mugal, C. F., Bolívar, P., Wang, M., Nadachowska-Brzyska, K., Smeds, L., Papoli, H., Gustafsson, L., & Ellegren, H. (2018). Sex-biased gene expression, sexual antagonism and levels of genetic diversity in the collared flycatcher (*Ficedula albicollis*) genome. Molecular Ecology, (March), 3572–3581. https://doi.org/10.1111/mec.14789

Eckhardt, F., Lewin, J., Cortese, R., Rakyan, V. K., Attwood, J., Burger, M., Burton, J., Cox, T. V, Davies, R., Down, T. A., Haefliger, C., Horton, R., Howe, K., … Beck, S. (2006). DNA methylation profiling of human chromosomes 6, 20 and 22. Nature Genetics, 38(12), 1378–1385. https://doi.org/10.1038/ng1909

Elgvin, T. O., Trier, C. N., Tørresen, O. K., Hagen, I. J., Lien, S., Nederbragt, A. J., Ravinet, M., Jensen, H., & Sætre, G.-P. (2017). The genomic mosaicism of hybrid speciation. Science Advances, 3(6), e1602996. https://doi.org/10.1126/sciadv.1602996

Ellegren, H., & Parsch, J. (2007). The evolution of sex-biased genes and sex-biased gene expression. Nature Reviews Genetics, 8(9), 689–698. https://doi.org/10.1038/nrg2167

Elmer, J. L., Hay, A. D., Kessler, N. J., Bertozzi, T. M., Ainscough, E. A. C., & Ferguson-Smith, A. C. (2021). Genomic properties of variably methylated retrotransposons in mouse. Mobile DNA, 12(1), 1–16. https://doi.org/10.1186/s13100-021-00235-1

Farrell, C., Thompson, M., Tosevska, A., Oyetunde, A., & Pellegrini, M. (2021). BiSulfite Bolt: A bisulfite sequencing analysis platform. GigaScience, 10(5), 1–8. https://doi.org/10.1093/gigascience/giab033

Fuentes, N., & Silveyra, P. (2019). Androgen Receptor Signaling. Cancer Research, 116, 135–170. https://doi.org/10.1158/0008-5472.can-03-3486

Gao, A., Su, J., Liu, R., Zhao, S., Li, W., Xu, X., Li, D., Shi, J., Gu, B., Zhang, J., Li, Q., Wang, X., Zhang, Y., … Wang, W. (2021). Sexual dimorphism in glucose metabolism is shaped by androgen-driven gut microbiome. Nature Communications, 12(1), 7080. https://doi.org/10.1038/s41467-021-27187-7

Girardet, L., Augière, C., Asselin, M. P., & Belleannée, C. (2019). Primary cilia: biosensors of the male reproductive tract. Andrology, 7(5), 588–602. https://doi.org/10.1111/andr.12650

Groen, K., Lea, R. A., Maltby, V. E., Scott, R. J., & Lechner-Scott, J. (2018). Letter to the editor: Blood processing and sample storage have negligible effects on methylation. Clinical Epigenetics, 10(1), 1–5. https://doi.org/10.1186/s13148-018-0455-6

Groothuis, T. G. G., & Schwabl, H. (2008). Hormone-mediated maternal effects in birds: Mechanisms matter but what do we know of them? Philosophical Transactions of the Royal Society B: Biological Sciences, 363(1497), 1647–1661. https://doi.org/10.1098/rstb.2007.0007

Gross, D. N., Van Den Heuvel, A. P. J., & Birnbaum, M. J. (2008). The role of FoxO in the regulation of metabolism. Oncogene, 27(16), 2320–2336. https://doi.org/10.1038/onc.2008.25

Gu, H., Smith, Z. D., Bock, C., Boyle, P., Gnirke, A., & Meissner, A. (2011). Preparation of reduced representation bisulfite sequencing libraries for genome-scale DNA methylation profiling. Nature Protocols, 6(4), 468–481. https://doi.org/10.1038/nprot.2010.190

Hall, E., Volkov, P., Dayeh, T., Esguerra, J. L. o. S., Salö, S., Eliasson, L., Rönn, T., Bacos, K., & Ling, C. (2014). Sex differences in the genome-wide DNA methylation pattern and impact on gene expression, microRNA levels and insulin secretion in human pancreatic islets. Genome Biology, 15(12), 522. https://doi.org/10.1186/s13059-014-0522-z

Hansen, K. D., Langmead, B., & Irizarry, R. A. (2012). BSmooth: from whole genome bisulfite sequencing reads to differentially methylated regions. Genome Biology, 13(10), R83. https://doi.org/10.1186/gb-2012-13-10-r83

Harrison, P. W., Wright, A. E., Zimmer, F., Dean, R., Montgomery, S. H., Pointer, M. A., & Mank, J. E. (2015). Sexual selection drives evolution and rapid turnover of male gene expression. Proceedings of the National Academy of Sciences, 112(14), 4393–4398. https://doi.org/10.1073/pnas.1501339112

Hartman, S., Taleb, S. A., Geng, T., Gyenai, K., Guan, X., & Smith, E. (2006). Comparison of plasma uric acid levels in five varieties of the domestic turkey, *Meleagris gallopavo*. Poultry Science, 85(10), 1791–1794. https://doi.org/10.1093/ps/85.10.1791

Head, J. A. (2014). Patterns of DNA methylation in animals: An ecotoxicological perspective. Integrative and Comparative Biology, 54(1), 77–86. https://doi.org/10.1093/icb/icu025

Hillier, L. W., Miller, W., Birney, E., Warren, W., Hardison, R. C., Ponting, C. P., Bork, P., Burt, D. W., Groenen, M. A. M., Delany, M. E., Dodgson, J. B., Chinwalla, A. T., Cliften, P. F., … Wilson, R. K. (2004). Sequence and comparative analysis of the chicken genome provide unique perspectives on vertebrate evolution. Nature, 432(7018), 695–716. https://doi.org/10.1038/nature03154

Hu, J., & Barrett, R. D. H. (2017). Epigenetics in natural animal populations. Journal of Evolutionary Biology, 30(9), 1612–1632. https://doi.org/10.1111/jeb.13130

Huang, L.-H., Lin, P.-H., Tsai, K.-W., Wang, L.-J., Huang, Y.-H., Kuo, H.-C., & Li, S.-C. (2017). The effects of storage temperature and duration of blood samples on DNA and RNA qualities. PLOS ONE, 12(9), e0184692. https://doi.org/10.1371/journal.pone.0184692

Husby, A. (2020). On the use of blood samples for measuring DNA methylation in ecological epigenetic studies. Integrative and Comparative Biology, 60(6), 1558–1566. https://doi.org/10.1093/icb/icaa123

Husby, A. (2022). Wild epigenetics : insights from epigenetic studies on natural populations. Proc. R. Soc. B, 289(1968), 2021.1633. https://doi.org/10.1098/rspb.2021.1633

Itoh, Y., Melamed, E., Yang, X., Kampf, K., Wang, S., Yehya, N., Van Nas, A., Replogle, K., Band, M. R., Clayton, D. F., Schadt, E. E., Lusis, A. J., & Arnold, A. P. (2007). Dosage compensation is less effective in birds than in mammals. Journal of Biology, 6(1). https://doi.org/10.1186/jbiol53

Jean-François, L., Victor, R., Morgane, T., Dominique, A., Vérane, B., Aurélie, C., Fernando, C. A. C. D., Michael, G., András, L. B. M. G. A., Alexander, S., Tamás, S., & Jean-Michel, G. (2020). Sex differences in adult lifespan and aging rates of mortality across wild mammals. Proceedings of the National Academy of Sciences, 117(15), 8546–8553. https://doi.org/10.1073/pnas.1911999117

Jones, P. A. (1999). The DNA methylation paradox. Trends in Genetics. https://doi.org/10.1016/S0168-9525(98)01636-9

Jones, P. A. (2012). Functions of DNA methylation: Islands, start sites, gene bodies and beyond. Nature Reviews Genetics, 13(7), 484–492. https://doi.org/10.1038/nrg3230

Kaiser, V. B., & Ellegren, H. (2006). Nonrandom distribution of genes with sex-biased expression in the chicken genome, 60(9), 1945–1951.

Laine, V. N., Gossmann, T. I., Schachtschneider, K. M., Garroway, C. J., Madsen, O., Verhoeven, K. J. F., De Jager, V., Megens, H. J., Warren, W. C., Minx, P., Crooijmans, R. P. M. A., Corcoran, P., Adriaensen, F., … Groenen, M. A. M. (2016). Evolutionary signals of selection on cognition from the great tit genome and methylome. Nature Communications, 7, 1–9. https://doi.org/10.1038/ncomms10474

Laine, V. N., Verhagen, I., Mateman, A. C., Pijl, A., Williams, T. D., Gienapp, P., Van Oers, K., & Visser, M. E. (2019). Exploration of tissue-specific gene expression patterns underlying timing of breeding in contrasting temperature environments in a song bird. BMC Genomics, 20(1), 1–16. https://doi.org/10.1186/s12864-019-6043-0

Langfelder, P., & Horvath, S. (2008). WGCNA: An R package for weighted correlation network analysis. BMC Bioinformatics, 9. https://doi.org/10.1186/1471-2105-9-559

Lawrence, M., Huber, W., Pagès, H., Aboyoun, P., Carlson, M., Gentleman, R., Morgan, M. T., & Carey, V. J. (2013). Software for computing and annotating genomic ranges. PLoS Computational Biology, 9(8), e1003118. https://doi.org/10.1371/journal.pcbi.1003118

Lea, A. J., Vilgalys, T. P., Durst, P. A. P., & Tung, J. (2017). Maximizing ecological and evolutionary insight in bisulfite sequencing data sets. Nature Ecology & Evolution, 1(8), 1074–1083. https://doi.org/10.1038/s41559-017-0229-0

Lewis, S., Benvenuti, S., Dall–Antonia, L., Griffiths, R., Money, L., Sherratt, T. N., Wanless, S., & Hamer, K. C. (2002). Sex-specific foraging behaviour in a monomorphic seabird. Proceedings of the Royal Society of London. Series B: Biological Sciences, 269(1501), 1687–1693. https://doi.org/10.1098/rspb.2002.2083

Li, M. W. M., Mruk, D. D., & Cheng, Y. C. (2009). Mitogen-activated protein kinases in male reproductive function. Trends in Molecular Medicine, 15(4), 159–168. https://doi.org/10.1016/j.molmed.2009.02.002

Li, Q., Li, N., Hu, X., Li, J., Du, Z., Chen, L., Yin, G., Duan, J., Zhang, H., Zhao, Y., Wang, J., & Li, N. (2011). Genome-wide mapping of DNA methylation in chicken. PLoS ONE, 6(5), 1–7. https://doi.org/10.1371/journal.pone.0019428

Li, Y., Pan, X., Roberts, M. L., Liu, P., Kotchen, T. A., Cowley, A. W., Mattson, D. L., Liu, Y., Liang, M., & Kidambi, S. (2018). Stability of global methylation profiles of whole blood and extracted DNA under different storage durations and conditions. Epigenomics, 10(6), 797–811. https://doi.org/10.2217/epi-2018-0025

Lindner, M., Verhagen, I., Viitaniemi, H. M., Laine, V. N., Visser, M. E., Husby, A., & van Oers, K. (2021). Temporal changes in DNA methylation and RNA expression in a small song bird: within- and between-tissue comparisons. BMC Genomics, 22(1), 1–16. https://doi.org/10.1186/s12864-020-07329-9

Loirand, G., & Pacaud, P. (2014). Small GTPases Involvement of Rho GTPases and their regulators in the pathogenesis of hypertension Involvement of Rho GTPases and their regulators in the pathogenesis of hypertension. Small GTPases, 5, e28846. https://doi.org/10.4161/sgtp.28846

Love, M. I., Huber, W., & Anders, S. (2014). Moderated estimation of fold change and dispersion for RNA-seq data with DESeq2. Genome Biology, 15(12), 1–21. https://doi.org/10.1186/s13059-014-0550-8

Luo, G. Z., Hafner, M., Shi, Z., Brown, M., Feng, G. H., Tuschl, T., Wang, X. J., & Li, X. C. (2012). Genome-wide annotation and analysis of zebra finch microRNA repertoire reveal sex-biased expression. BMC Genomics, 13(1). https://doi.org/10.1186/1471-2164-13-727

Luo, R., Bai, C., Yang, L., Zheng, Z., Su, G., Gao, G., Wei, Z., Zuo, Y., & Li, G. (2018). Correction to: DNA methylation subpatterns at distinct regulatory regions in human early embryos. Open Biology, 8(12), 1–9. https://doi.org/10.1098/rsob.180215

Mank, J. E., Hultin-Rosenberg, L., Webster, M. T., & Ellegren, H. (2008). The unique genomic properties of sex-biased genes: Insights from avian microarray data. BMC Genomics, 9, 1–14. https://doi.org/10.1186/1471-2164-9-148

Mannucci, A., Argento, F. R., Fini, E., Coccia, M. E., Taddei, N., Becatti, M., & Fiorillo, C. (2022). The Impact of oxidative stress in male infertility. Frontiers in Molecular Biosciences, 8(January), 1–9. https://doi.org/10.3389/fmolb.2021.799294

Maunakea, A. K., Nagarajan, R. P., Bilenky, M., Ballinger, T. J., Dsouza, C., Fouse, S. D., Johnson, B. E., Hong, C., Nielsen, C., Zhao, Y., Turecki, G., Delaney, A., Varhol, R., … Costello, J. F. (2010). Conserved role of intragenic DNA methylation in regulating alternative promoters. Nature, 466(7303), 253–257. https://doi.org/10.1038/nature09165

McCarthy, N. S., Melton, P. E., Cadby, G., Yazar, S., Franchina, M., Moses, E. K., Mackey, D. A., & Hewitt, A. W. (2014). Meta-analysis of human methylation data for evidence of sex-specific autosomal patterns. BMC Genomics, 15(1). https://doi.org/10.1186/1471-2164-15-981

McCormick, H., Young, P. E., Hur, S. S. J., Booher, K., Chung, H., Cropley, J. E., Giannoulatou, E., & Suter, C. M. (2017). Isogenic mice exhibit sexually-dimorphic DNA methylation patterns across multiple tissues. BMC Genomics, 18(1), 966. https://doi.org/10.1186/s12864-017-4350-x

Mishra, I., Sharma, A., Prabhat, A., Batra, T., Malik, I., & Kumar, V. (2020). Changes in the expression of genes involved in DNA methylation and histone modification in response to daily food availability times in zebra finches: Epigenetic implications. Journal of Experimental Biology, 223(3). https://doi.org/10.1242/jeb.217422

Moghadam, H., Pointer, M., Wright, A., Sofia, B., & Mank, J. (2012). W chromosome expression responds to female-specific selection. Proceedings of the National Academy of Sciences, 109(21), 8207–8211. https://doi.org/10.1073/pnas.1202721109

Mugal, C. F., Wang, M., Backström, N., Wheatcroft, D., Ålund, M., Sémon, M., McFarlane, S. E., Dutoit, L., Qvarnström, A., & Ellegren, H. (2020). Tissue-specific patterns of regulatory changes underlying gene expression differences among *Ficedula* flycatchers and their naturally occurring F1 hybrids. Genome Research, 31(12), 1727–1739. https://doi.org/10.1101/gr.254508.119

Nätt, D., Agnvall, B., & Jensen, P. (2014). Large sex differences in chicken behaviour and brain gene expression coincide with few differences in promoter DNA-methylation. PLoS ONE, 9(4). https://doi.org/10.1371/journal.pone.0096376

Nätt, D., Rubin, C.-J., Wright, D., Johnsson, M., Beltéky, J., Andersson, L., & Jensen, P. (2012). Heritable genome-wide variation of gene expression and promoter methylation between wild and domesticated chickens. BMC Genomics, 13(1), 59. https://doi.org/10.1186/1471-2164-13-59

Naurin, S., Hansson, B., Hasselquist, D., Kim, Y. H., & Bensch, S. (2011). The sex-biased brain: Sexual dimorphism in gene expression in two species of songbirds. BMC Genomics, 12. https://doi.org/10.1186/1471-2164-12-37

Naurin, S., Hasselquist, D., Bensch, S., & Hansson, B. (2012). Sex-biased gene expression on the avian Z chromosome: Highly expressed genes show higher male-biased Expression. PLoS ONE, 7(10). https://doi.org/10.1371/journal.pone.0046854

Nie, H., Crooijmans, R. P. M. A., Lammers, A., van Schothorst, E. M., Keijer, J., Neerincx, P. B. T., Leunissen, J. A. M., Megens, H. J., & Groenen, M. A. M. (2010). Gene expression in chicken reveals correlation with structural genomic features and conserved patterns of transcription in the terrestrial vertebrates. PLoS ONE, 5(8), 1–7. https://doi.org/10.1371/journal.pone.0011990

Ortiz-Huidobro, R. I., Velasco, M., Larqué, C., Escalona, R., & Hiriart, M. (2021). Molecular insulin actions are sexually dimorphic in lipid metabolism. Frontiers in Endocrinology. https://doi.org/10.3389/fendo.2021.690484

Pabbidi, M. R., Kuppusamy, M., Didion, S. P., Sanapureddy, P., Reed, J. T., & Sontakke, S. P. (2018). Sex differences in the vascular function and related mechanisms: role of 17β-estradiol. American Journal of Physiology-Heart and Circulatory Physiology, 315(6), H1499–H1518. https://doi.org/10.1152/ajpheart.00194.2018

Pellman, B. A., Schuessler, B. P., Tellakat, M., & Kim, J. J. (2017). Sexually dimorphic risk mitigation strategies in rats. Eneuro, 4(1), ENEURO.0288-16.2017. https://doi.org/10.1523/ENEURO.0288-16.2017

Peona, V., Palacios-Gimenez, O. M., Blommaert, J., Liu, J., Haryoko, T., Jønsson, K. A., Irestedt, M., Zhou, Q., Jern, P., & Suh, A. (2021). The avian W chromosome is a refugium for endogenous retroviruses with likely effects on female-biased mutational load and genetic incompatibilities. Philosophical Transactions of the Royal Society B: Biological Sciences, 376(1833). https://doi.org/10.1098/rstb.2020.0186

Piao, Y., Xu, W., Park, K. H., Ryu, K. H., & Xiang, R. (2021). Comprehensive evaluation of differential methylation analysis methods for bisulfite sequencing data. International Journal of Environmental Research and Public Health, 18(15), 7975. https://doi.org/10.3390/ijerph18157975

Platt 2nd, R. N., Blanco-Berdugo, L., & Ray, D. A. (2016). Accurate transposable element annotation is vital when analyzing new genome assemblies. Genome Biology and Evolution, 8(2), 403–410. https://doi.org/10.1093/gbe/evw009

Qi, L. M., Mohr, M., & Wade, J. (2012). Enhanced expression of tubulin-specific chaperone protein a, mitochondrial ribosomal protein S27, and the DNA excision repair protein XPACCH in the song system of juvenile male zebra finches. Developmental Neurobiology, 72(2), 199–207. https://doi.org/10.1002/dneu.20956

Quinlan, A. R., & Hall, I. M. (2010). BEDTools: A flexible suite of utilities for comparing genomic features. Bioinformatics, 26(6), 841–842. https://doi.org/10.1093/bioinformatics/btq033

Richani, D., & Gilchrist, R. B. (2018). The epidermal growth factor network: Role in oocyte growth, maturation and developmental competence. Human Reproduction Update, 24(1), 1–14. https://doi.org/10.1093/humupd/dmx029

Rinn, J. L., & Snyder, M. (2005). Sexual dimorphism in mammalian gene expression. Trends in Genetics. https://doi.org/10.1016/j.tig.2005.03.005

Rizzo, A., Roscino, M. T., Binetti, F., & Sciorsci, R. L. (2012). Roles of reactive oxygen species in female reproduction. Reproduction in Domestic Animals, 47(2), 344–352. https://doi.org/10.1111/j.1439-0531.2011.01891.x

Rønning, B., Broggi, J., Bech, C., Børge, M., Ringsby, T. H., Parn, H., Hagen, I. J., Sæther, B. E., & Jensen, H. (2016). Is basal metabolic rate associated with recruit production and survival in free-living house sparrows? Functional Ecology, 30, 1140–1148. https://doi.org/10.1111/1365-2435.12597

Rubenstein, D. R., Skolnik, H., Berrio, A., Champagne, F. A., Phelps, S., & Solomon, J. (2016). Sex-specific fitness effects of unpredictable early life conditions are associated with DNA methylation in the avian glucocorticoid receptor. Molecular Ecology, 25(8), 1714–1728. https://doi.org/10.1111/mec.13483

Sæther, S., Sætre, G.-P., Borge, T., Wiley, C., Svedin, N., Andersson, G., Veen, T., Haavie, J., Servedio, M., Bureš, S., Král, M., Mårten, H., Gustafsson, L., … Qvarnström, A. (2007). Sex chromosome-linked species recognition and evolution of reproductive isolation in flycatchers. Science, 318(5847), 95–97. https://doi.org/10.1126/science.1141506

Sætre, G., Borge, T., Lindroos, K., Haavie, J., Sheldon, B. C., Primmer, C., & Syvänen, A. (2003). Sex chromosome evolution and speciation in *Ficedula* flycatchers. Proceedings of the Royal Society of London. Series B: Biological Sciences, 270(1510), 53–59. https://doi.org/10.1098/rspb.2002.2204

San Agustin, J. T., Pazour, G. J., & Witman, G. B. (2015). Intraflagellar transport is essential for mammalian spermiogenesis but is absent in mature sperm. Molecular Biology of the Cell, 26(24), 4358–4372. https://doi.org/10.1091/mbc.E15-08-0578

Sargent, K. M., McFee, R. M., Gomes, R. S., & Cupp, A. S. (2015). VEGFA: Just one of multiple mechanisms for sex-specific vascular development within the testis? Journal of Endocrinology, 227(2), R31–R50. https://doi.org/10.1530/JOE-15-0342

Schröder, C., & Steimer, W. (2018). gDNA extraction yield and methylation status of blood samples are affected by long-term storage conditions. PloS One, 13(2), e0192414–e0192414. https://doi.org/10.1371/journal.pone.0192414

Seale, L. A., Ogawa-Wong, A. N., & Berry, M. J. (2018). Sexual dimorphism in selenium metabolism and selenoproteins. Free Radical Biology and Medicine, 127, 198–205. https://doi.org/10.1016/j.freeradbiomed.2018.03.036

Sharma, E., Künstner, A., Fraser, B. A., Zipprich, G., Kottler, V. A., Henz, S. R., Weigel, D., & Dreyer, C. (2014). Transcriptome assemblies for studying sex-biased gene expression in the guppy, *Poecilia reticulata*. BMC Genomics, 15(1), 400. https://doi.org/10.1186/1471-2164-15-400

Sheleg, M., Yochum, C., Richardson, J., Wagner, G., & Zhou, R. (2015). Ephrin-a5 regulates inter-male aggression in mice. Behav Brain Res, 286, 300–307. https://doi.org/10.1016/j.bbr.2015.03.001

Själander, M., Jahre, M., Tufte, G., & Reissmann, N. (2019). EPIC: An energy-efficient, high-performance GPGPU computing research infrastructure. ArXiv. https://doi.org/10.48550/arxiv.1912.05848

Sudmant, P., Alexis, M., & Burge, C. (2015). Meta-analysis of RNA-seq expression data across species, tissues and studies. Genome Biology, 16, 287. https://doi.org/10.1186/s13059-015-0853-4

Tang, Y. P., & Wade, J. (2010). Sex- and age-related differences in ribosomal proteins L17 and L37, as well as androgen receptor protein, in the song control system of zebra finches. Neuroscience, 171(4), 1131–1140. https://doi.org/10.1016/j.neuroscience.2010.10.014

Tarka, M., Guenther, A., Niemelä, P. T., Nakagawa, S., & Noble, D. W. A. (2018). Sex differences in life history, behavior, and physiology along a slow-fast continuum: a meta-analysis. Behavioral Ecology and Sociobiology, 72(8). https://doi.org/10.1007/s00265-018-2534-2

Uebbing, S., Künstner, A., Mäkinen, H., Backström, N., Bolivar, P., Burri, R., Dutoit, L., Mugal, C. F., Nater, A., Aken, B., Flicek, P., Martin, F. J., Searle, S. M. J., & Ellegren, H. (2016). Divergence in gene expression within and between two closely related flycatcher species. Molecular Ecology, 25(9), 2015–2028. https://doi.org/10.1111/mec.13596

Uebbing, S., Künstner, A., Mäkinen, H., & Ellegren, H. (2013). Transcriptome sequencing reveals the character of incomplete dosage compensation across multiple tissues in flycatchers. Genome Biology and Evolution, 5(8), 1555–1566. https://doi.org/10.1093/gbe/evt114

Valdebenito, J. O., Halimubieke, N., Lendvai, Á. Z., Figuerola, J., Eichhorn, G., & Székely, T. (2021). Seasonal variation in sex-specific immunity in wild birds. Scientific Reports, 11(1), 1–11. https://doi.org/10.1038/s41598-020-80030-9

Viitaniemi, H. M., Verhagen, I., Visser, M. E., Honkela, A., Van Oers, K., Husby, A., & Meyer, M. (2019). Seasonal variation in genome-wide DNA methylation patterns and the onset of seasonal timing of reproduction in great tits. Genome Biology and Evolution, 11(3), 970–983. https://doi.org/10.1093/gbe/evz044

Vincze, O., Vágási, C. I., Pénzes, J., Szabó, K., Magonyi, N. M., Czirják, G., & Pap, P. L. (2022). Sexual dimorphism in immune function and oxidative physiology across birds: The role of sexual selection. Ecology Letters, 25(4), 958–970. https://doi.org/10.1111/ele.13973

Watson, H., Powell, D., Salmón, P., Jacobs, A., & Isaksson, C. (2020). Urbanization is associated with modifications in DNA methylation in a small passerine bird. Evolutionary Applications, 14(1), 85–98. https://doi.org/10.1111/eva.13160

Wen, W., Cho, Y. S., Zheng, W., Dorajoo, R., Kato, N., Qi, L., Chen, C. H., Delahanty, R. J., Okada, Y., Tabara, Y., Gu, D., Zhu, D., Haiman, C. A., … Shu, X. O. (2012). Meta-analysis identifies common variants associated with body mass index in east Asians. Nature Genetics, 44(3), 307–311. https://doi.org/10.1038/ng.1087

Wu, X., Zhang, Q., Xu, S., Jin, P., Luan, P., Li, Y., Cao, Z., Leng, L., Wang, Y., & Wang, S. (2016). Differential expression of six chicken genes associated with fatness traits in a divergently selected broiler population. Molecular and Cellular Probes, 30(1), 1–5. https://doi.org/10.1016/j.mcp.2015.12.003

Xu, Q., Yuan, Z., Dongxiao, S., Yachun, W., & Ying, Y. (2007). Analysis on DNA methylation of various tissues in chicken. Animal Biotechnology, 18, 231–241. https://doi.org/10.1080/10495390701574838

Yan, X., Liu, H., Liu, J., Zhang, R., Wang, G., Li, Q., Wang, D., Li, L., & Wang, J. (2015). Evidence in duck for supporting alteration of incubation temperature may have influence on methylation of genomic DNA. Poultry Science, 94, 2537–2545. https://doi.org/10.3382/ps/pev201

