## Supplementary information for "DNA methylation regulates sex-biased gene expression in the house sparrow"

**Table S1:** Statistics for raw sequence data. Individual ID is shown alongside mappability and mean percentage methylation for RRBS data, and total sequences and mappability for RNA-seq data. Information for males is shown on page 1, and information for females follows on page 2.

| ID | Tissue | RRBS mappability % | RRBS methylation % | RNAseq sequence count | RNAseq read count |
|---|---|---|---|---|---|
| **8M71323** | Blood | 86.56 | 24.12 | - | - |
| | Brain | 87.90 | 32.12 | 49079718 | 8349020 |
| | Liver | 85.74 | 26.83 | 52198172 | 12009638 |
| | Testes | 85.95 | 26.06 | 48311739 | 9539250 |
| **8N06837** | Blood | 85.34 | 27.78 | - | - |
| | Brain | 85.07 | 26.97 | 51023683 | 9195781 |
| | Liver | 86.51 | 27.74 | 62704138 | 15616217 |
| | Testes | 85.55 | 24.93 | 56685281 | 12133232 |
| **8N72694** | Blood | 86.47 | 21.01 | - | - |
| | Brain | 87.24 | 32.30 | 60229663 | 11459631 |
| | Liver | 85.87 | 27.31 | 64494078 | 14679664 |
| | Testes | 86.64 | 24.18 | 49530661 | 9570022 |
| **8N73488** | Blood | 86.66 | 24.76 | - | - |
| | Brain | 87.90 | 34.31 | 45488840 | 7891619 |
| | Liver | 86.98 | 28.02 | 54496681 | 11770184 |
| | Testes | 85.22 | 23.92 | 23880042 | 4427567 |
| **8N73907** | Blood | 85.46 | 28.27 | - | - |
| | Brain | 86.89 | 28.91 | 49126940 | 8591262 |
| | Liver | 84.80 | 27.82 | 88407816 | 18097925 |
| | Testes | 86.30 | 28.62 | 29155321 | 6282298 |
| **8N87513** | Blood | 84.89 | 27.05 | - | - |
| | Brain | 87.63 | 34.31 | 48163735 | 8067713 |
| | Liver | 86.67 | 26.42 | 71658239 | 15769044 |
| | Testes | 86.37 | 24.06 | 46675446 | 9703642 |
| **8N87539** | Blood | 86.08 | 21.16 | - | - |
| | Brain | 84.78 | 26.13 | 58843238 | 10666312 |
| | Liver | 86.11 | 28.89 | 68623166 | 17403528 |
| | Testes | 87.31 | 26.76 | 58426570 | 13080386 |
| **8N87590** | Blood | 85.30 | 23.86 | - | - |
| | Brain | 87.07 | 33.20 | 61024352 | 9860038 |
| | Liver | 85.48 | 25.52 | 49281167 | 10917064 |
| | Testes | 84.81 | 25.40 | 55743893 | 11587042 |

| | | | | | |
|---|---|---|---|---|---|
| **8N72676** | Blood | 85.21 | 28.81 | - | - |
| | Brain | 86.48 | 34.74 | 57763713 | 9835204 |
| | Liver | 85.94 | 27.81 | 67970782 | 17338838 |
| | Ovary | 86.86 | 28.42 | 51661731 | 9900298 |
| **8N73920** | Blood | 85.98 | 24.10 | - | - |
| | Brain | 85.88 | 28.85 | 56018919 | 9303743 |
| | Liver | 85.08 | 26.60 | 56241301 | 12171345 |
| | Ovary | 85.94 | 25.11 | 45673592 | 10435744 |
| **8N87534** | Blood | 86.35 | 24.76 | - | - |
| | Brain | 87.30 | 33.64 | 48504503 | 7752887 |
| | Liver | 85.83 | 26.03 | 52817980 | 11456236 |
| | Ovary | 86.73 | 28.78 | 49952070 | 10044279 |
| **8N87536** | Blood | 86.10 | 24.01 | - | - |
| | Brain | 85.05 | 26.09 | 45872466 | 7933132 |
| | Liver | 85.51 | 27.27 | 61052080 | 12244259 |
| | Ovary | 85.02 | 28.25 | 61661833 | 13907944 |
| **8N87543** | Blood | 84.10 | 19.27 | - | - |
| | Brain | 86.49 | 31.13 | 47846085 | 8730748 |
| | Liver | 85.92 | 27.75 | 37554715 | 9096325 |
| | Ovary | 85.78 | 25.46 | 57651910 | 13414334 |
| **8N87582** | Blood | 86.60 | 26.39 | - | - |
| | Brain | 86.67 | 30.96 | 53792872 | 9564822 |
| | Liver | 86.31 | 26.66 | 66222201 | 14094965 |
| | Ovary | 84.43 | 26.81 | 56914251 | 12496539 |
| **8N87594** | Blood | 85.88 | 23.76 | - | - |
| | Brain | 87.58 | 34.46 | 55001603 | 9110788 |
| | Liver | 86.27 | 29.63 | 72406170 | 14883073 |
| | Ovary | 85.99 | 28.40 | 62671298 | 13340787 |
| **8N87595** | Blood | 85.24 | 20.56 | - | - |
| | Brain | 86.83 | 33.34 | 59380888 | 10304596 |
| | Liver | 85.52 | 25.90 | 54431565 | 11746813 |
| | Ovary | 85.67 | 25.50 | 46876005 | 10599329 |
| | **MEAN:** | **86.06** | **27.31** | **54774857** | **11174481** |

**Table S2:** Number of sites with different read depths and mean methylation percentage, first when given read depth is shared for all samples between tissues (n = 16 individuals, n = 64 samples), then when given read depth is shared for all samples within each tissue (n = 16 individuals).

| | No filtering | 3x | 5x | 10x |
|---|---|---|---|---|
| **All tissues (n= 16 individuals, n = 64 samples)** | | | | |
| CpG sites | 3380542 | 561846 | **313701** | 76367 |
| Mean methylation % | 27.853 | 14.190 | **13.498** | 13.207 |
| **Blood (n = 16)** | | | | |
| CpG sites | 2607065 | 786402 | **533416** | 173610 |
| Mean methylation % | 26.066 | 13.107 | **11.843** | 11.214 |
| **Brain (n = 16)** | | | | |
| CpG sites | 3104436 | 938690 | **644517** | 263211 |
| Mean methylation % | 30.392 | 27.191 | **28.528** | 30.828 |
| **Liver (n = 16)** | | | | |
| CpG sites | 3184622 | 1111914 | **852190** | 368452 |
| Mean methylation % | 26.633 | 22.765 | **22.661** | 23.648 |
| **Gonad (n = 16)** | | | | |
| CpG sites | 3109628 | 905552 | **585205** | 167454 |
| Mean methylation % | 25.344 | 20.770 | **20.768** | 22.585 |

**Table S3:** Separate file. Overview of differentially expressed genes that were identified for each tissue in DEG analysis using DeSeq2.

**Table S4:** Separate file. Results of functional analysis of DMRs identified in differential methylated region analyses in Methylkit, and of functional analysis of DEGs identified in differential gene expression analysis using DeSeq2. For blood (DMGs only), brain, and liver we used all sex-biased genes detected for functional analysis due to low number of identified genes. For gonad we used only the genes that had DMRs in their TSS or promoter and that were also DEGs. To further reduce gonad associated functional terms to the most relevant terms, "real hub genes" (the 5% of genes in the network with highest degree connectivity) were selected before functional analysis using StringApp. Functional analysis was performed using GO Biological Process, Reactome Pathway, Wiki Pathway, KEGG pathway, and TISSUE annotations. A redundancy cutoff of 0.6 was used to collapse redundant terms.

**Table S5:** Separate file. Transcripts belonging to WGCNA modules.

**Table S6:** Separate file. Results of functional analysis for the “real hub” genes from WGCNA modules that were identified from PPI networks produced using the Cytoscape StringApp. For each WGCNA module that was strongly correlated with sex (the turquoise, pink, and black modules) hub genes were first identified (module membership > 0.8, p.GS.sex < 0.05) and used to produce a STRING PPI network using the default edge confidence of 0.4. Subsequently, “real hub” genes were identified for each network using a connectivity degree that selected the most connected ~5% of nodes in each network. Functional analysis was performed using GO Biological Process, Reactome Pathway, Wiki Pathway, KEGG pathway, and TISSUE annotations. A redundancy cutoff of 0.6 was used to collapse redundant terms.

**Table S7:** Separate file. Overview of differentially methylated genes (DMGs) with DMRs in the TSS or promoter that were identified for each tissue in DMR analyses using Methylkit.

**Table S8:** FDR corrected *p*-values from comparison of mean methylation values between different genomic features (10 Kbp upstream of annotated gene start, promoter, TSS, gene body, as well as transposable elements CR1s and LTRs) using pairwise Wilcoxon rank sum tests. Methylation levels differed significantly between all genomic features, but were most similar between promoters and CR1s.

| REGION | + 10 Kbp | Promoter | TSS | Gene body | CR1s |
|---|---|---|---|---|---|
| **Promoter** | $5.811e^{-173}$ | - | - | - | - |
| **TSS** | $< 1e^{-308}$ | $5.408e^{-142}$ | - | - | - |
| **Gene body** | $7.109e^{-180}$ | $< 1e^{-308}$ | $< 1e^{-308}$ | - | - |
| **CR1s** | $1.483e^{-70}$ | $1.381e^{-4}$ | $7.518e^{-29}$ | $3.102e^{-179}$ | - |
| **LTRs** | $1.927e^{-218}$ | $< 1e^{-308}$ | $< 1e^{-308}$ | $3.252e^{-55}$ | $3.458e^{-244}$ |

**Table S9:** Spearman's correlation between mean DNA methylation percentage and rld transformed expression values for different genomic features in brain, liver, and gonad. Two-sided tests were used for all features.

| Feature | Estimate | *p*-value | Method |
|---|---|---|---|
| **Brain** | | | |
| 10kb upstream | -0.143 | $2.783e^{-20}$ | Spearman's rank correlation rho |
| Gene body | -0.079 | $1.877e^{-7}$ | Spearman's rank correlation rho |
| Promoter | -0.274 | $3.093e^{-63}$ | Spearman's rank correlation rho |
| TSS | -0.366 | $1.669e^{-62}$ | Spearman's rank correlation rho |
| **Liver** | | | |
| 10kb upstream | -0.083 | $1.585e^{-7}$ | Spearman's rank correlation rho |
| Gene body | -0.124 | $2.956e^{-16}$ | Spearman's rank correlation rho |
| Promoter | -0.225 | $7.225e^{-42}$ | Spearman's rank correlation rho |
| TSS | -0.372 | $7.249e^{-68}$ | Spearman's rank correlation rho |
| **Gonad** | | | |
| 10kb upstream | -0.163 | $4.530e^{-26}$ | Spearman's rank correlation rho |
| Gene body | -0.122 | $3.294e^{-16}$ | Spearman's rank correlation rho |
| Promoter | -0.326 | $7.587e^{-91}$ | Spearman's rank correlation rho |
| TSS | -0.449 | $5.261e^{-94}$ | Spearman's rank correlation rho |

**Table S10:** Overlap between sex-biased differentially expressed genes (DEGs) and differentially methylated genes (DMGs) in brain, liver, and gonad. Here we look at only the DEGs that had at least one 300 bp region containing at least five CpG sites in their TSS or promoter in brain (30%), liver (33%), and gonad (38%). In gonad 14% of DEGs were also DMGs, but 48% of DMGs were also DEGs. Conversely, very few genes were both differentially methylated and differentially expressed in brain and liver.

| | Brain DMGs | Liver DMGs | Gonad DMGs |
|---|---|---|---|
| **Brain DEGs** | **2 (2%)** | 0 (0%) | 33 (2%) |
| **Liver DEGs** | 0 (0%) | **0 (0%)** | 26 (1%) |
| **Gonad DEGs** | 7 (8%) | 1 (7%) | **364 (14%)** |

### Figures

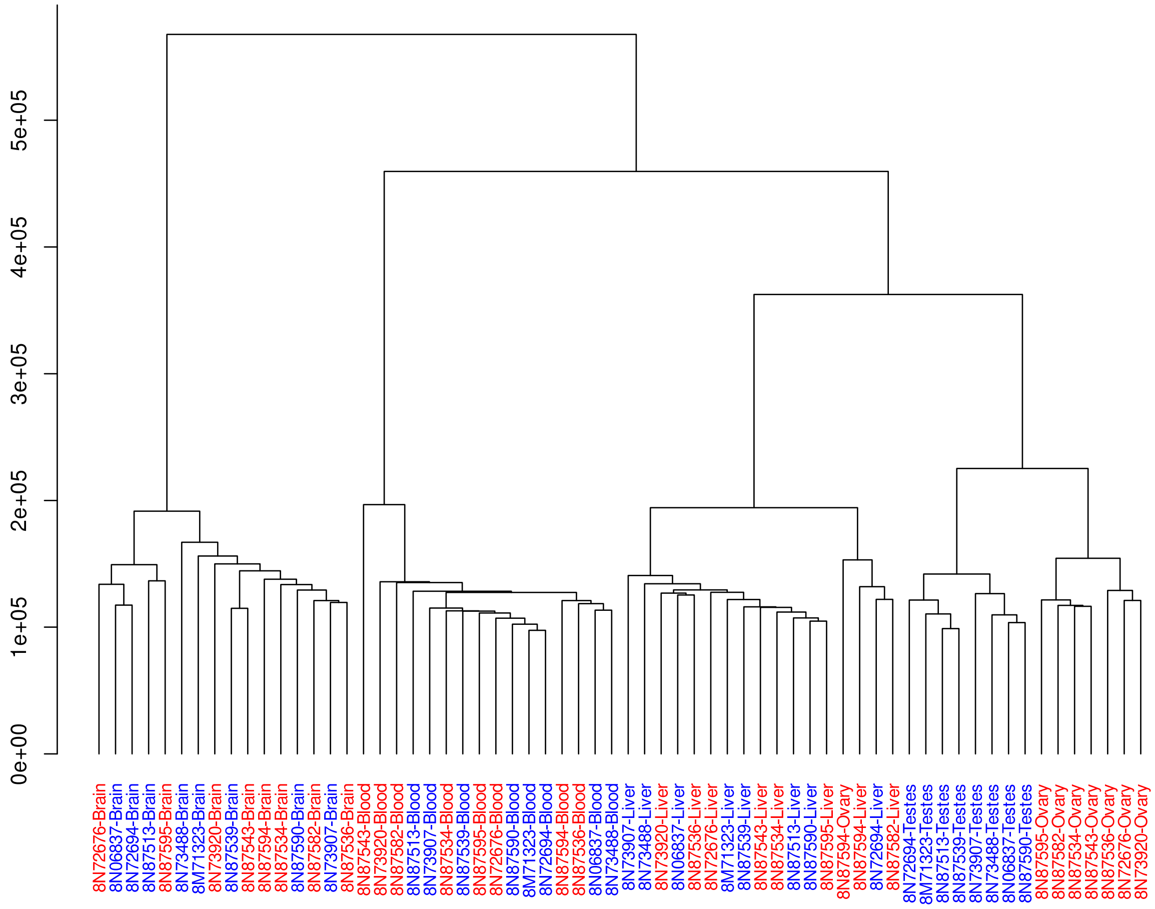


**Figure S1:** Cluster dendrogram for sites with 5x coverage shared between all samples across all tissues (313701 sites). Hierarchical Ward D2 clustering with Manhattan distance was used. Samples from females are shown in red, and samples from males in blue. Sample names include individual ring number, as well as tissue from which the DNA used for bisulfite sequencing was extracted. Samples clustered by sex in the gonad, but clustering by sex was not observed for blood, brain, or liver tissue.

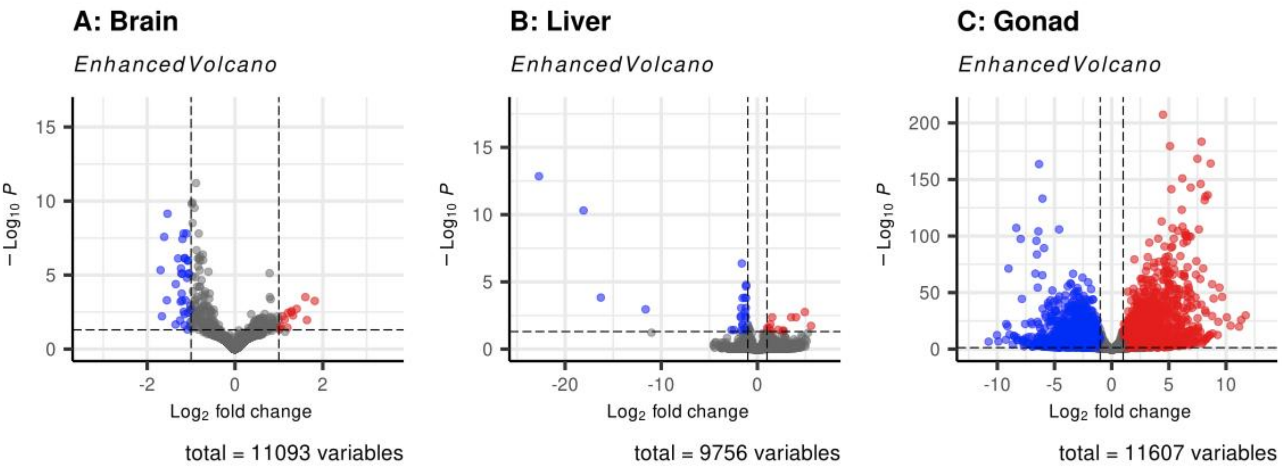


**Figure S2:** Volcano plots of all the transcripts for each tissue analysed in differential gene expression analysis using DeSeq2. Genes that were differentially expressed with $p < 0.05$ after correcting for false discovery rate are shown in red if they were more highly expressed in females, or in blue if they were more highly expressed in males (n = 300 DEGs for brain, n = 45 DEGs for liver, and n = 6838 DEGs for gonad). Genes with $p >= 0.05$ after correcting for false discovery rate are shown in grey.

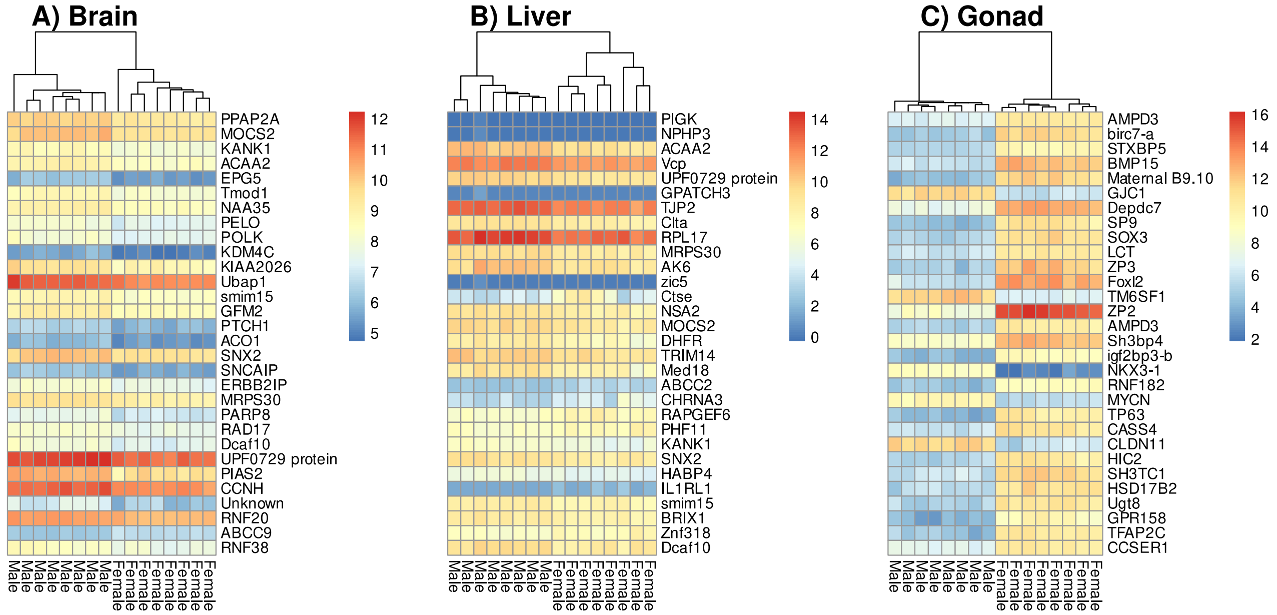


**Figure S3:** Heatmap of the top 30 sex-biased differentially expressed genes (DEGs) for each tissue from differential gene expression analyses using DeSeq2. Heatmaps are coloured according to rld regularised gene expression for each sample, and clustered according to sex at these genes. Note that the scale of the heatmap is different for each tissue, and that the greatest differences in gene expression between the sexes were observed in the gonad.

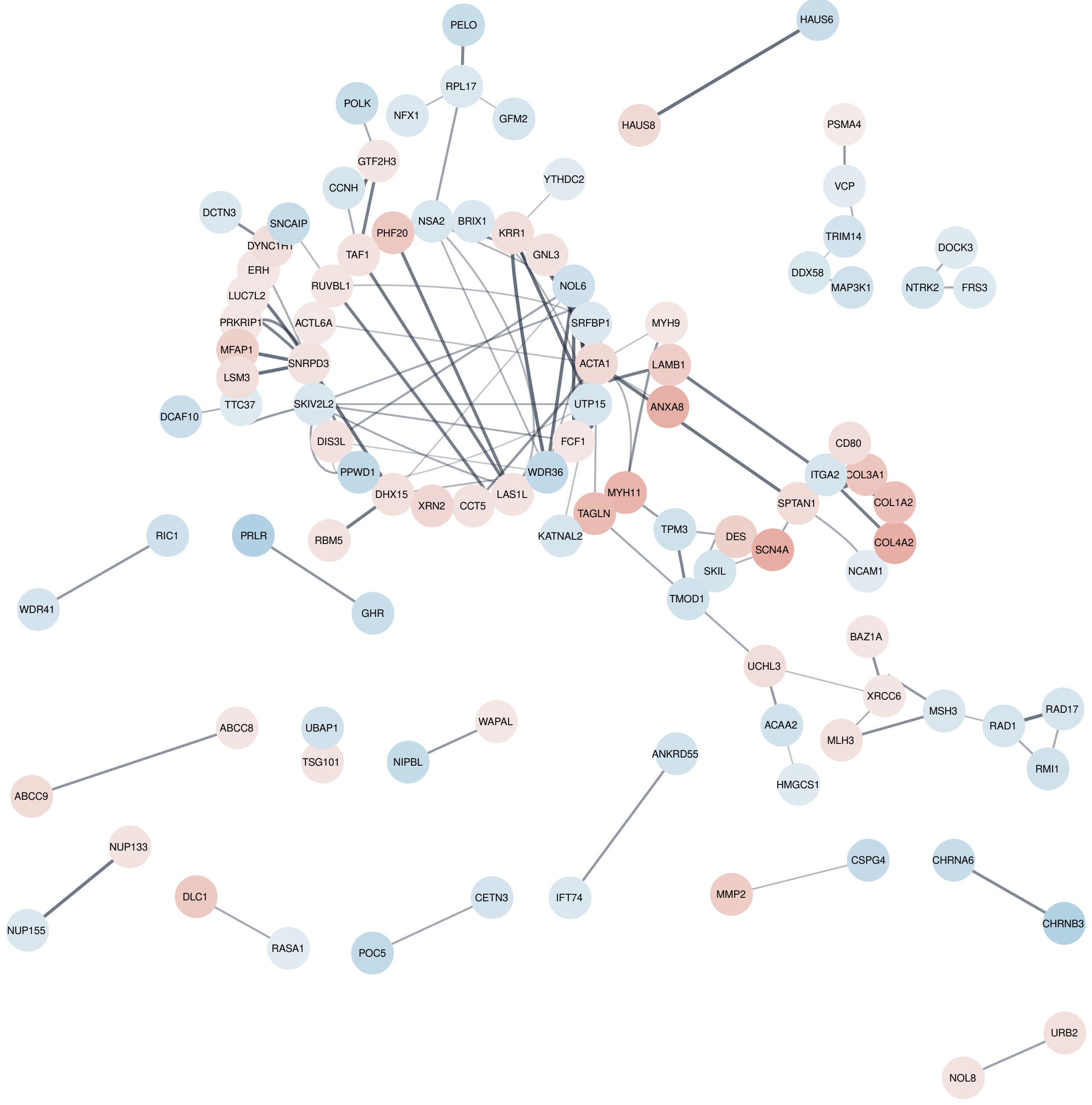


**Figure S4A:** PPI network of all differentially expressed genes (DEGs) for brain. The network was drawn using Cytoscape StringApp with a default edge confidence of 0.4. Nodes are coloured according to log2 fold change (positive values indicate genes that are more highly expressed in females and are coloured red, negative values indicate genes that are more highly expressed in males and are coloured blue). See Table S6 for a list of functional terms identified in functional analysis on these DEGs using STRING.

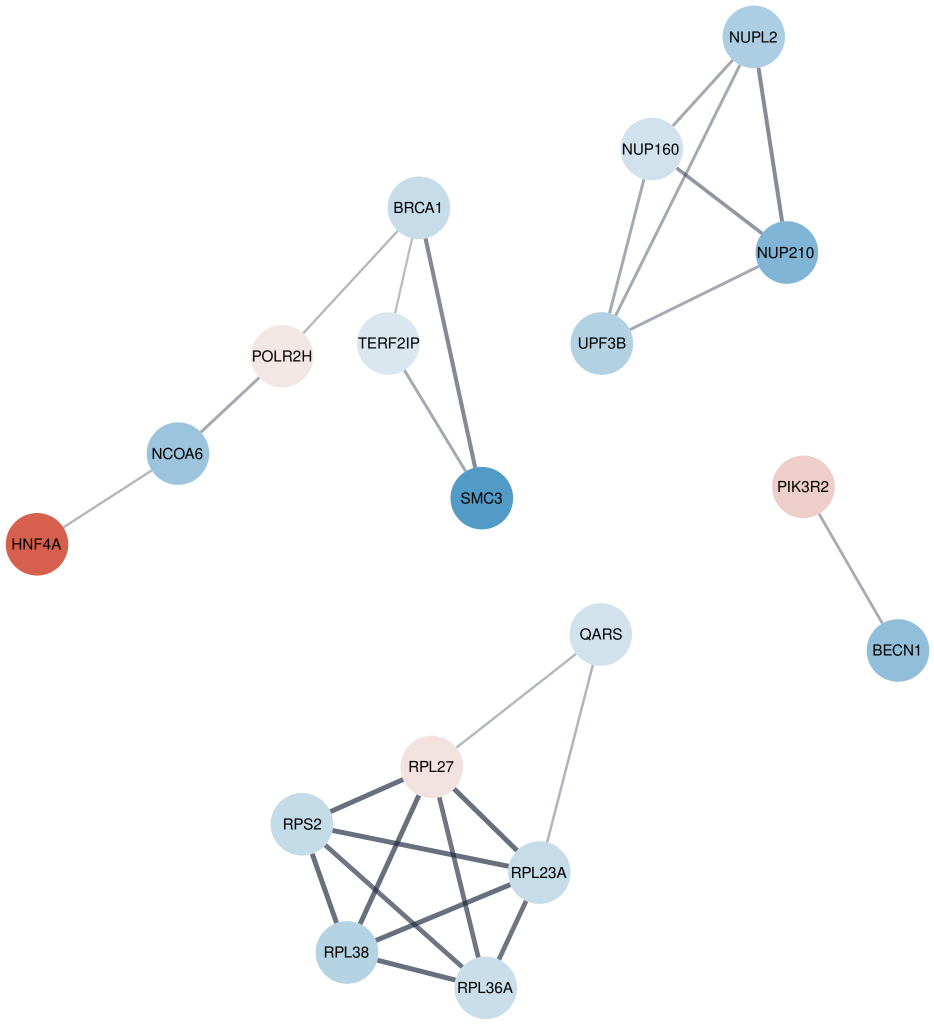


**Figure S4B:** PPI network of “real hub” genes for gonad, drawn using Cytoscape StringApp with a default edge confidence of 0.4. Nodes are coloured according to log2 fold change (positive values indicate genes that are more highly expressed in females and are coloured red, negative values indicate genes that are more highly expressed in males and are coloured blue). First a PPI network was drawn using the default edge confidence of 0.4, and using only the genes that had sex-biased DMRs in their TSS or promoter and that were also DEGs. The “real hub” genes shown here were then identified as the 5% of genes in the PPI network with highest degree connectivity, this was done to reduce the many gonad associated functional terms to the most relevant terms. See Table S6 for a list of functional terms identified in functional analysis on these “real hub” genes using STRING.

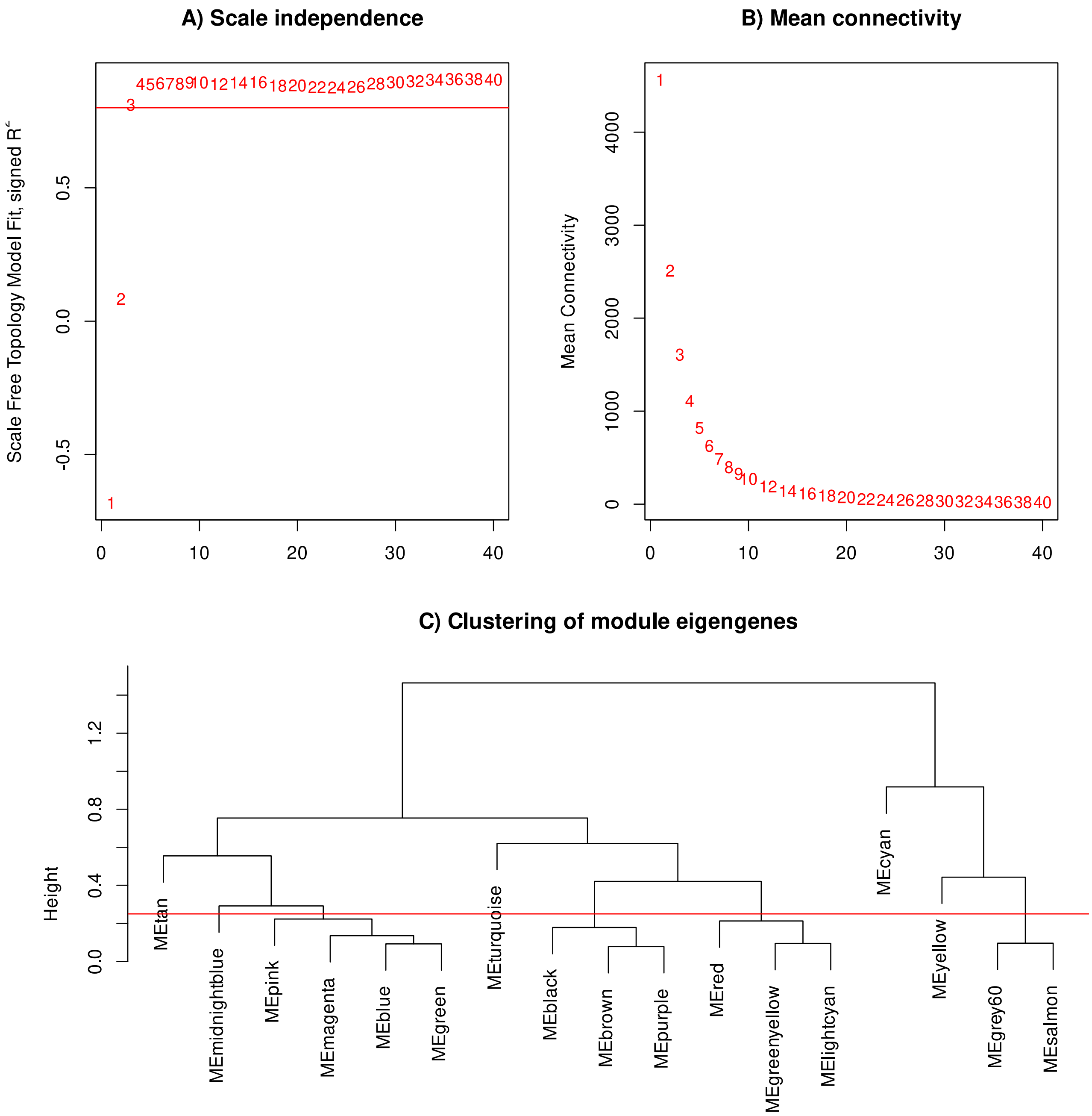


**Figure S5: A)** Scale independence (scale free topology fit index as a function of soft-thresholding power) and **B)** main connectivity for gonad normalised count data. Based on plots A and B, a soft-threshold power of 4 was chosen for subsequent WGCNA analysis because 4 is the lowest power for which the scale-free topology fit index reaches 0.8. **C)** To merge modules with similar expression profiles a height of 0.25 was chosen, which corresponds to correlation of 0.75.

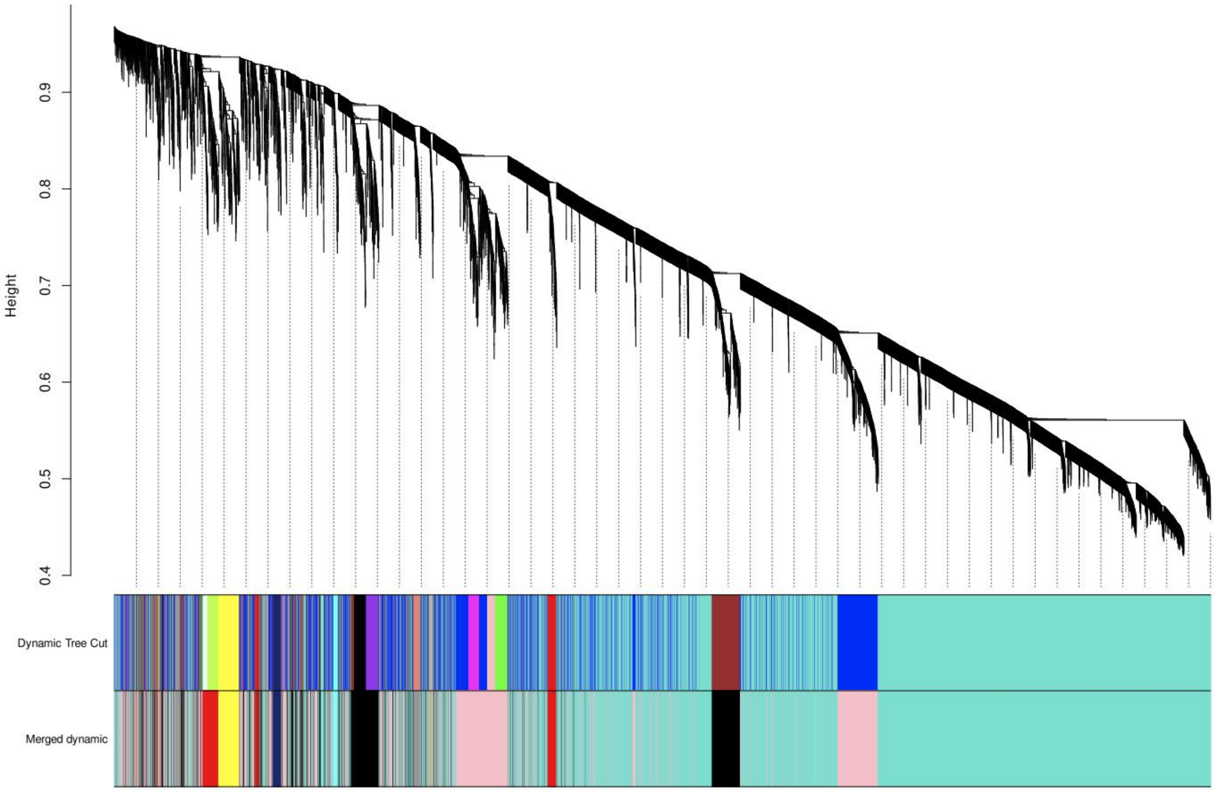


**Figure S6:** Clustering dendrogram for all modules based on WGCNA modules (dynamic tree cut) and WGCNA module eigengenes (merged dynamic), before and after merging modules for which the expression profiles were similar.

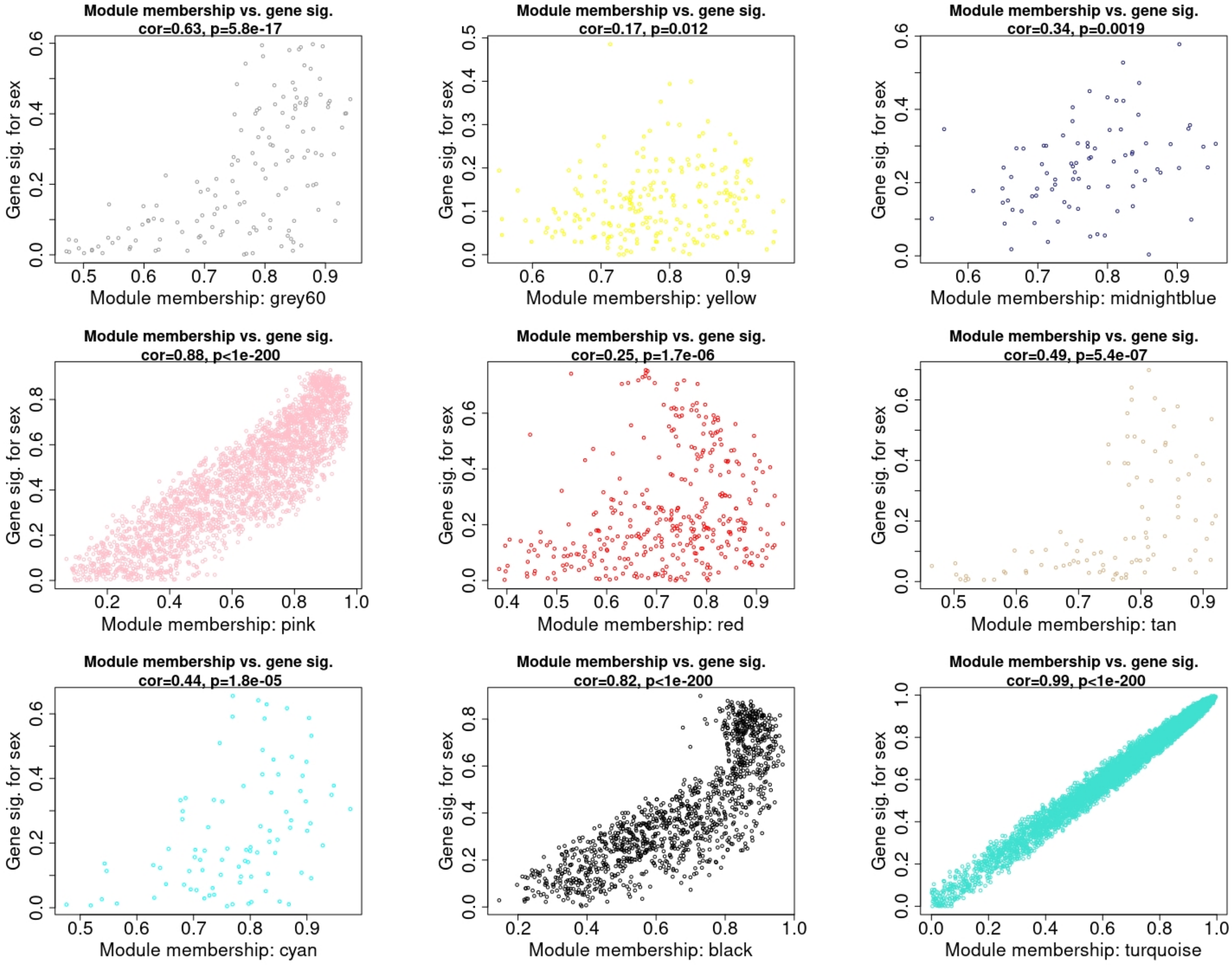


**Figure S7:** Module membership (x) versus gene significance for sex (y) for all genes present in each identified WGCNA co-expression module. Modules with correlation score of > 0.8 (the turquoise, pink, and black modules) were used to produce PPI networks of the hub genes from each module. Hub genes in each module were defined as those with module membership > 0.8 and $p < 0.05$ for the significance of the relationship with sex. For the turquoise module n = 3808 hub genes, for the pink module n = 761 hub genes, and for the black module n = 342 hub genes.

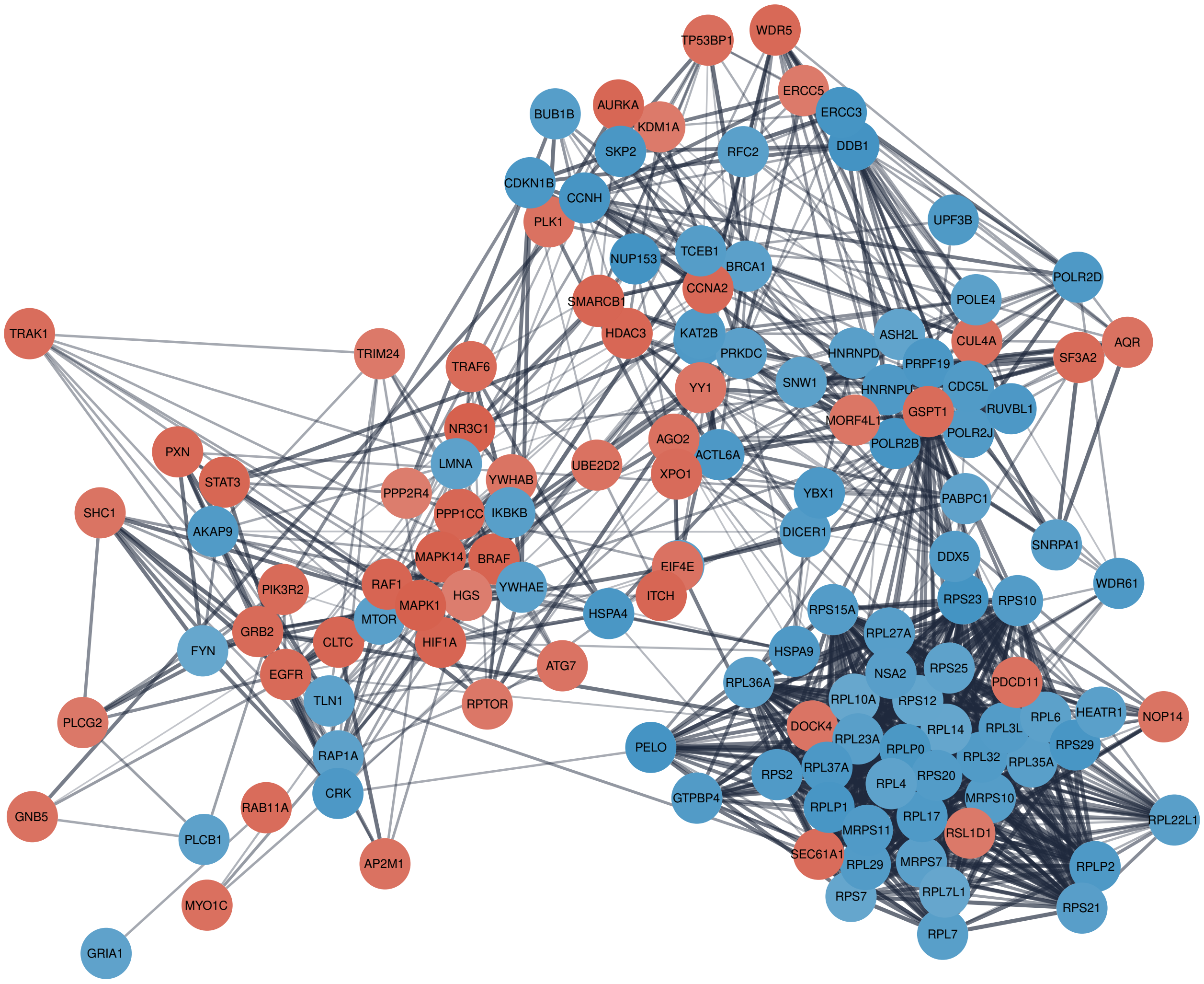


**Figure S8A:** The PPI network of “real hub” genes for the turquoise module that was positively correlated with sex. Real hub genes were defined as the ~5% of genes with greatest degree connectivity (in + out) in the full PPI network of hub genes. The edge thickness represents node connectivity where thicker lines indicate more evidence for the connection based on experimental evidence of protein-protein interaction. Nodes are coloured according to log2 fold change (positive values indicate genes that are more highly expressed in females and are coloured red, negative values indicate genes that are more highly expressed in males and are coloured blue).

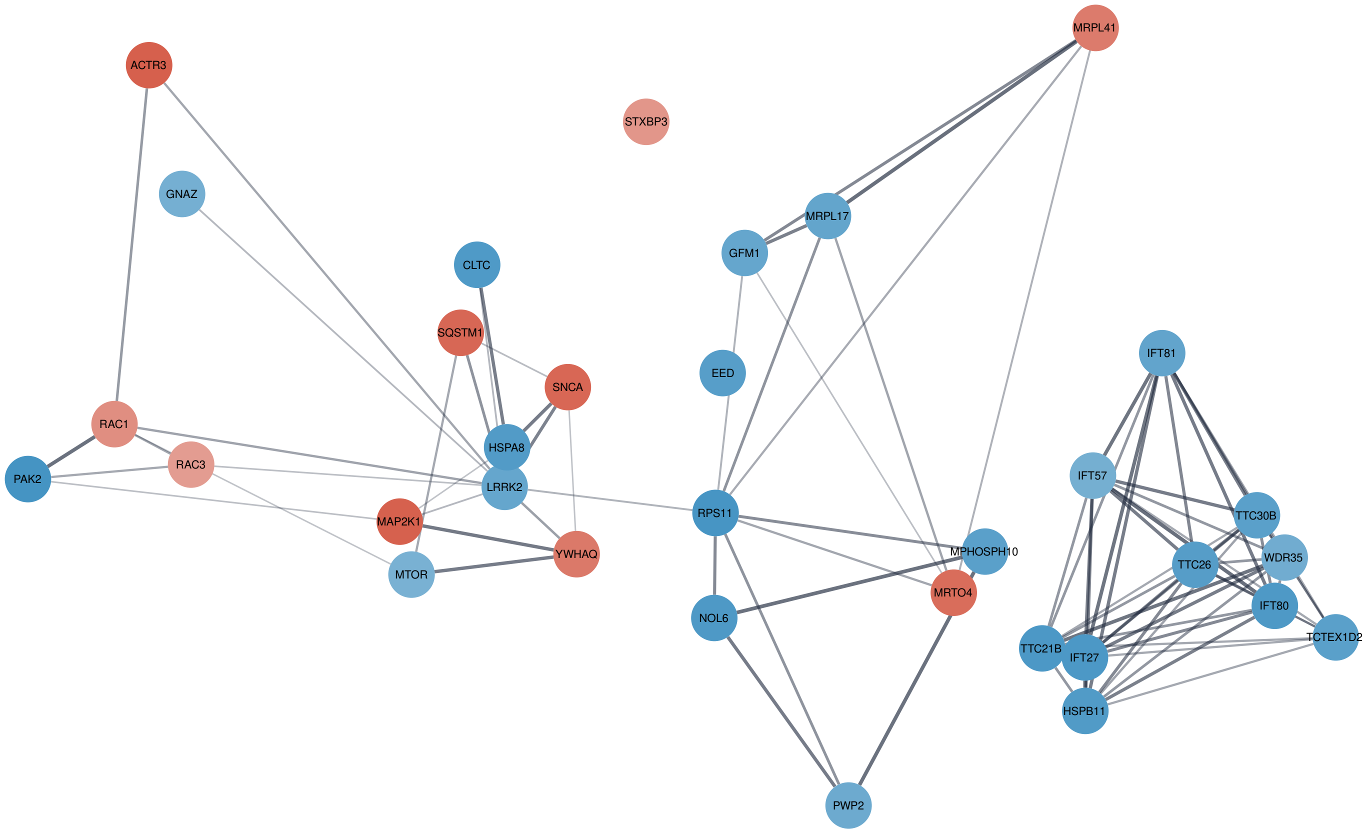


**Figure S8B:** The PPI network of “real hub” genes for the pink module that was negatively correlated with sex. Real hub genes were defined as the ~5% of genes with greatest degree connectivity (in + out) in the full PPI network of hub genes. The edge thickness represents node connectivity where thicker lines indicate more evidence for the connection based on experimental evidence of protein-protein interaction. Nodes are coloured according to log2 fold change (positive values indicate genes that are more highly expressed in females and are coloured red, negative values indicate genes that are more highly expressed in males and are coloured blue).

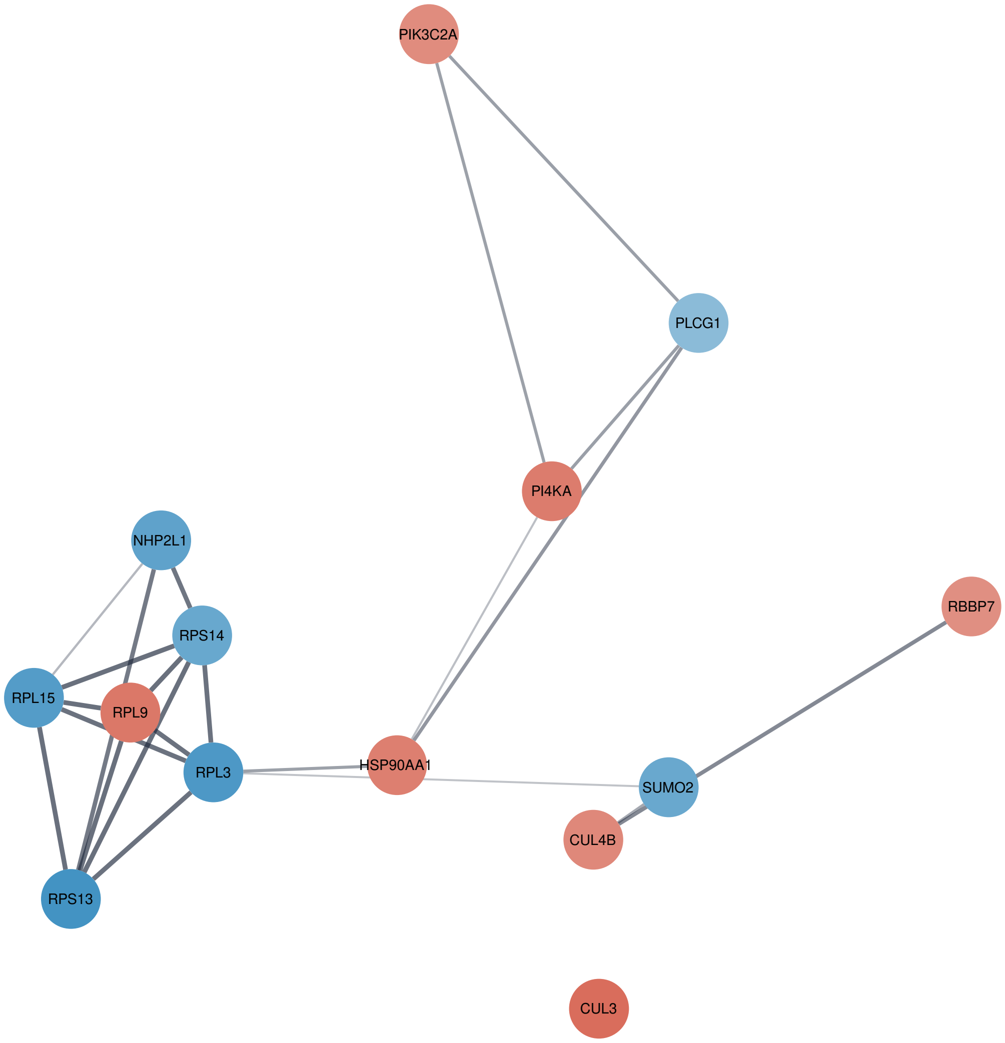


**Figure S8C:** The PPI network of "real hub" genes for the black module that was positively correlated with sex. Real hub genes were defined as the ~5% of genes with greatest degree connectivity (in + out) in the full PPI network of hub genes. The edge thickness represents node connectivity where thicker lines indicate more evidence for the connection based on experimental evidence of protein-protein interaction. Nodes are coloured according to log2 fold change (positive values indicate genes that are more highly expressed in females and are coloured red, negative values indicate genes that are more highly expressed in males and are coloured blue).

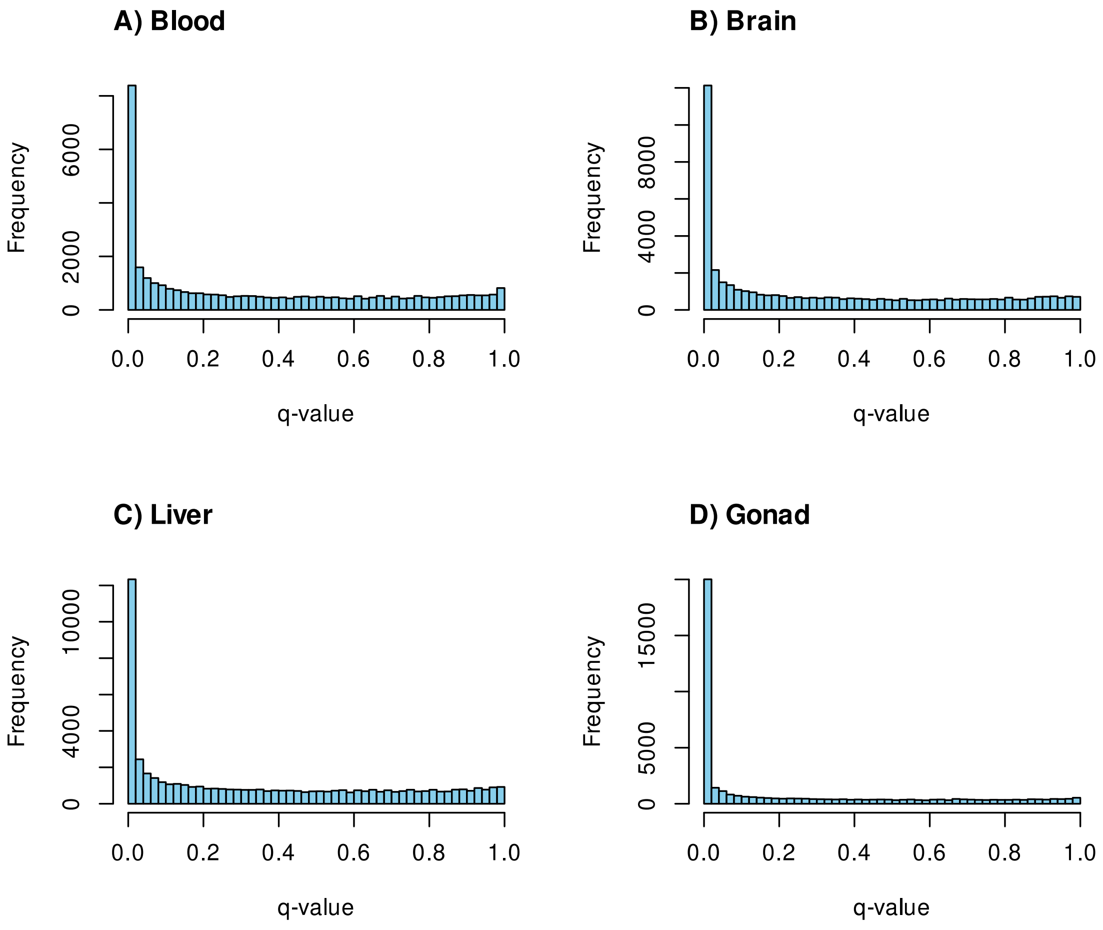


**Figure S9:** Raw *q*-value distributions from differential methylated region analysis using Methylkit, prior to DMR selection (DMRs were defined as sites with at least 10% methylation difference between males and females and *q*-value < 0.05). For DMR analysis the genome was tiled into windows of 300 bp and only tiles that contained at least 5 CpGs were used in subsequent DMR analyses. Selecting tiles in this manner eliminated the second peak at *q*-value = 1.0 that is often observed for differential methylation analysis on single CpG sites. This is because tiles containing at least 5 CpGs are more likely to show some variation in methylation level between test groups (here between sexes) than single CpGs, whereas in differential methylated site analyses there may be many sites that do not vary between test groups.

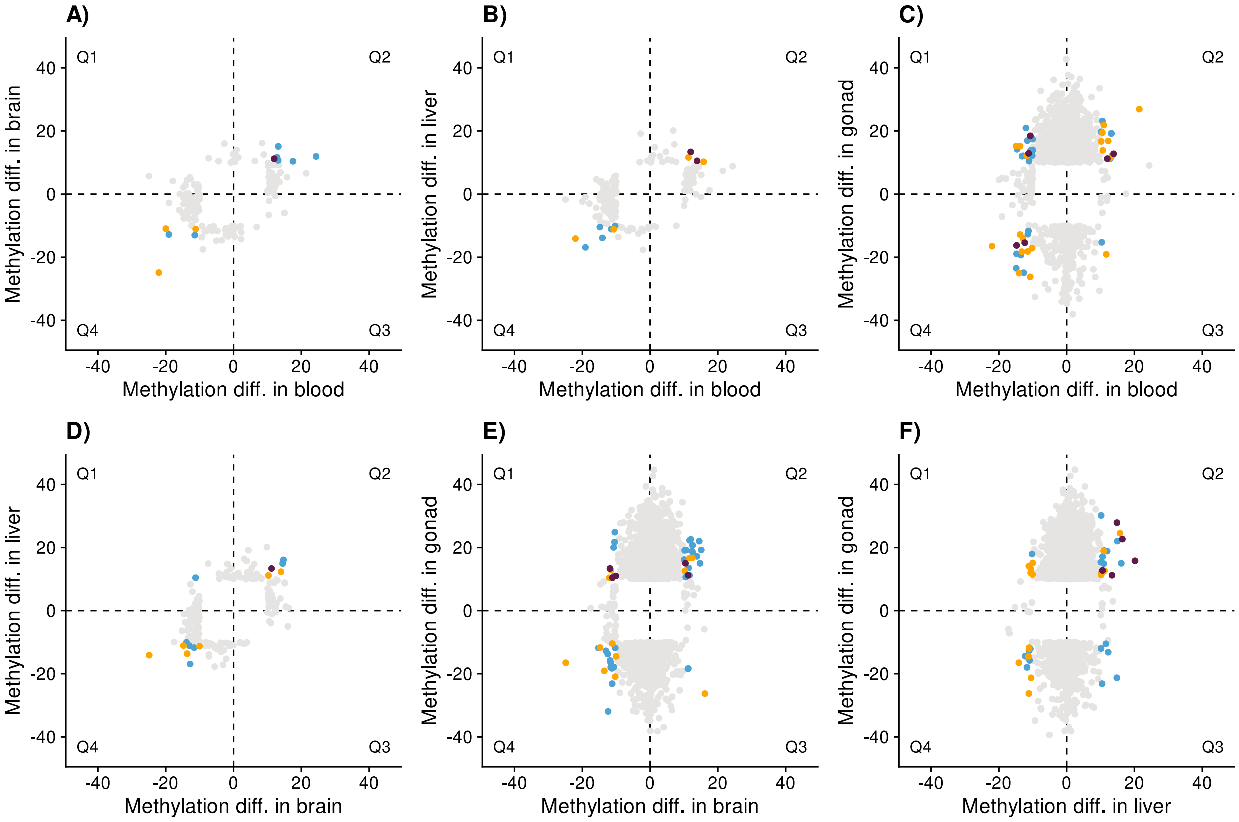


**Figure S10:** Quadrant plots showing tissue comparisons of the methylation difference between males and females at DMRs (when males are coded as 1 and females as 2). For DMRs that were detected in both tissues in a comparison, those in the TSS are shown in dark purple, those in the promoter are shown in orange, and those in the gene body are shown in blue. DMRs that were only detected in one tissue in a comparison are shown in grey. The 4 quadrants are separated by dotted lines and labelled Q1-Q4. Over representation of points in Q2 and Q4 is expected because we a-priori expect methylation levels at a given region to be similar between tissues. This pattern is not observed for tissue comparisons with gonad, thus methylation differences between the sexes at important regions in the gonad often do not correlate with methylation differences between the sexes at the same regions in other tissues. This underscores the importance of choosing the appropriate tissue for the analysed phenotype when carrying out studies on DNA methylation.

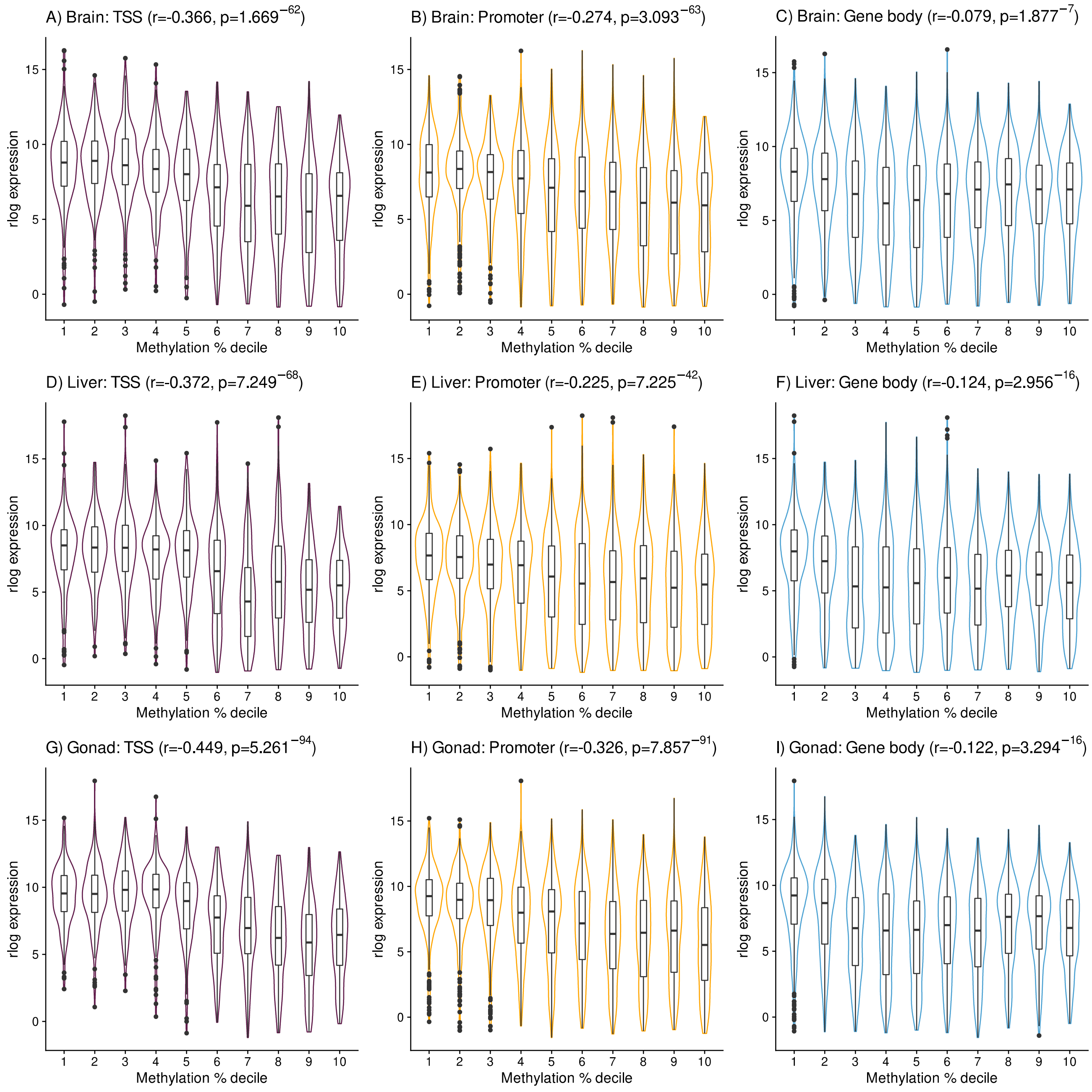


**Figure S11:** Violin plot of methylation percentage decile (x) against rld regularised gene expression values (y) for each genomic feature (TSS, promoter, gene body) for each tissue (brain, liver, gonad). Mean methylation percentage was first calculated for each genomic feature for each gene using 5x CpG sites shared between all samples within each tissue (for brain 644517 sites, for liver 852190 CpG sites, and for gonad 585205 CpG sites). Then mean methylation percentage at these genomic features was divided into deciles for plotting.

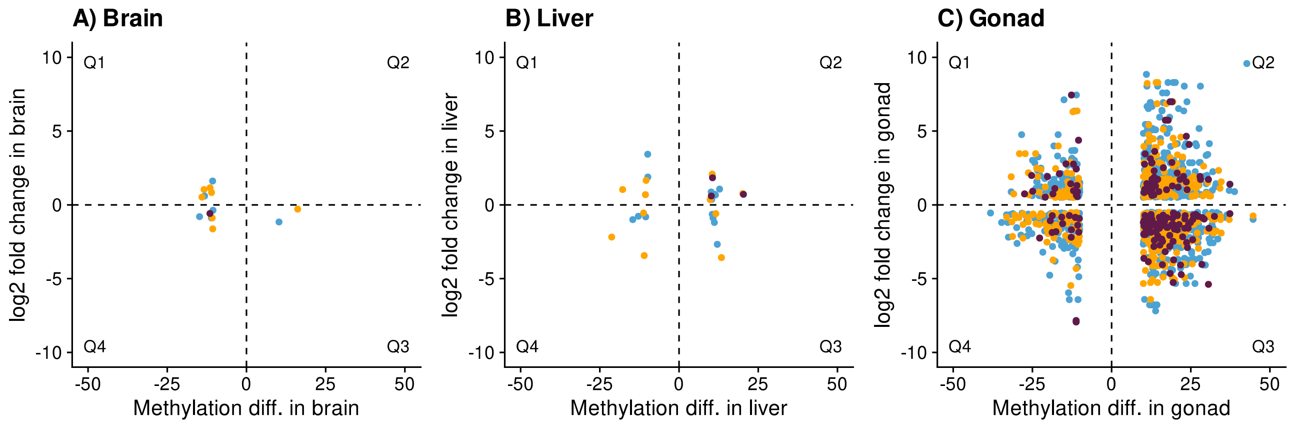


**Figure S12:** Quadrant plots showing methylation difference between males and females in relation to log2 fold change between males and females (when male = 1 and female = 2). DMRs in the promoter of genes are shown in orange, DMRs in the TSS are shown in dark purple, and DMRs in the gene body are shown in blue. The 4 quadrants are separated by dotted lines and labelled Q1-Q4. Over representation of points in Q1 and Q3 is expected for DMRs located in transcription start sites compared to points for DMRs in gene bodies, and this is what we observe for gonad (Fisher's Exact Test *p*-value = 0.021).
